# Comprehensive Characterization of BTK Inhibitor Specificity, Potency, and Biological Effects: Insights into Covalent and Non-covalent Mechanistic Signatures

**DOI:** 10.1101/2024.09.06.611550

**Authors:** Antonia C. Darragh, Andrew M. Hanna, Justin H. Lipner, Nicole B. Servant, Alastair J. King, Mirza Jahic

## Abstract

Uncovering a drug’s mechanism of action and possible adverse effects are critical components in drug discovery and development. Moreover, it provides evidence for why some drugs prove more effective than others, and how to design better drugs altogether. Here we demonstrate the utility of a high-throughput *in vitro* screening platform along with a comprehensive panel to aid in the characterization of fifteen BTK inhibitors that are either approved by the FDA or presently under clinical evaluation. To compare the potency of these drugs, we measured the binding affinity of each to wild-type BTK, as well as a clinically relevant resistance mutant of BTK (BTK C481S). In doing so, we discovered a considerable difference in the selectivity and potency of these BTK inhibitors to the wild-type and mutant proteins. Some of this potentially contributes to the adverse effects experienced by patients undergoing therapy using these drugs. Overall, non-covalent BTK inhibitors showed stronger potency for both the wild-type and mutant BTK when compared with that of covalent inhibitors, with the majority demonstrating a higher specificity and less off-target modulation. Additionally, we compared biological outcomes for four of these inhibitors in human cell-based models. As expected, we found different phenotypic profiles for each inhibitor. However, the two non-covalent inhibitors had fewer off-target biological effects when compared with the two covalent inhibitors. This and similar in-depth preclinical characterization of drug candidates can provide critical insights into the efficacy and mechanism of action of a compound that may affect its safety in a clinical setting.

**Table of Contents/Abstract Graphic:** 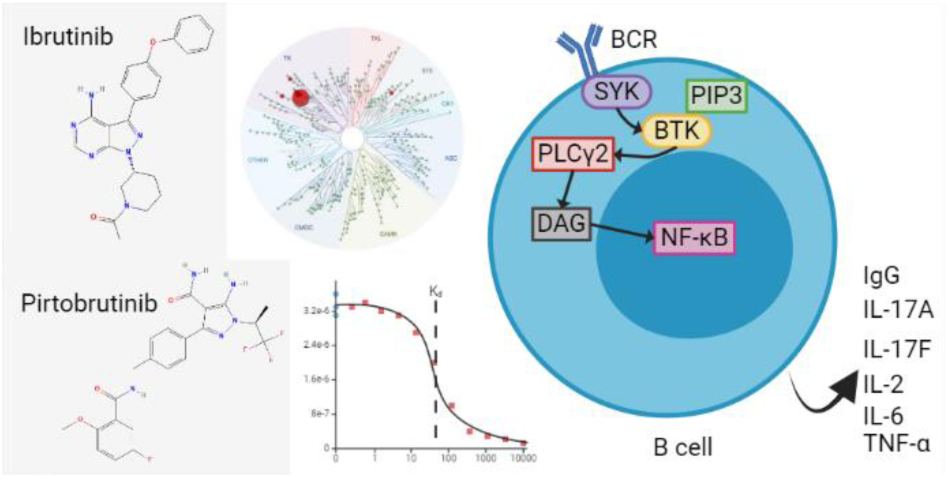

---

Kinases are important regulatory proteins that serve various signaling mechanisms *in vivo*^1^. A compromised kinase, for example through over/under-activation or a mutation, may result in the progression of various diseases such as cancer^1^ and for kinases that function in the immune system, autoimmune diseases^2^. Bruton’s tyrosine kinase (BTK) is a non-receptor protein-tyrosine kinase in the TEC family^3^. BTK is downstream of the B cell receptor (BCR) signaling cascade that promotes the development of various immune cell types (e.g., ^4^). BTK is critical for B cell development and the function of mature B cells, playing a significant role in B cell malignancies (e.g., ^5–7^). The first U.S. Food and Drug Administration (FDA)-approved BTK inhibitor, ibrutinib, was found to not only be efficacious for B cell malignancies but also autoimmune disease^8^. Ibrutinib has been very successful in treating patients; however, adverse effects of the drug have been observed (e.g., ^9^). For example, mild to moderate diarrhea, nausea, and fatigue^10, 11^ and atrial fibrillation^12^ have been observed following ibrutinib treatment. Fortunately, these adverse effects are considerably less severe than those from non-specific chemotherapy, which used to be the primary treatment for B cell malignancies^13^. However, adverse effects are much less tolerated when treating immune diseases as compared with cancer^13^.

Apart from adverse events, a more common reason for the discontinuation of ibrutinib treatment is the development of resistance, often through a cysteine to serine mutation in BTK (BTK C481S)^14–17^. Next-generation BTK inhibitors have focused on maintaining ibrutinib’s B cell malignancy efficacy and/or the treatment of autoimmune diseases with fewer adverse effects and less resistance (e.g., ^18, 19^).

At least 27 BTK inhibitors have entered clinical trials, six of which have been approved for treatment in humans by at least one government agency^13^. Acalabrutinib became the first-in-class treatment for mantle cell lymphoma (MCL) by the FDA in 2017^20^. In 2023, zanubrutinib became the first-in-class treatment in the U.S. for chronic lymphocytic leukemia (CLL) and small lymphocytic lymphoma (SLL), and is now approved for the treatment of relapsed or refractory follicular lymphoma^20^. In Japan, tirabrutinib was approved by the Pharmaceuticals and Medical Devices Agency (PMDA) for the treatment of primary central nervous system diffuse large B cell lymphoma (PCNS DLBCL) in 2020^21, 22^. China’s National Medicines and Pharmaceutical Administration (NMPA) approved orelabrutinib for the treatment of MCL, CLL, and SLL in 2020^23^. Pirtobrutinib (LOXO-305) is the first non-covalent BTK inhibitor to be granted FDA approval^24^. It received accelerated approval in 2023 for the treatment of relapsed or refractory (R/R) MCL after treatment with at least one other BTK inhibitor and another systemic therapy; as well as CLL and SLL treatment after treatment with at least one other BTK inhibitor and a BCL2 inhibitor^24^. Pirtobrutinib treatment overcomes some BTK inhibitor resistance through its ability to bind to BTK mutants, including BTK C481S^16, 25^. Unfortunately, resistance to pirtobrutinib treatment occurs through the accumulation of other BTK mutations (e.g., at residues T474 and V416) and second-site kinase-dead mutations^16, 26^.

Most BTK inhibitors hinder BTK activity by binding to the active site of BTK, i.e., its ATP-binding pocket, usually at the same residue (C481)^26, 27^ (Figure 1). While most BTK inhibitors bind covalently (irreversibly) to this particular residue, some of the next-generation drugs include non-covalent inhibitors, some of which have been found to inhibit this mutated form of BTK^16^. In addition to resistance mutations that change this residue, 10 other kinases have a cysteine residue at an analogous location in their active site, making them likely off-targets of BTK inhibitors^28^. These kinases are the other four TEC family kinases (BMX, ITK, TEC, and TXK) and six other kinases (BLK, EGFR, ERBB2, ERBB4, JAK3, and MKK7)^29^. These off-targets are thought to contribute to some of the adverse effects of BTK inhibitor treatment^18, 30^. For example, inhibition of the tyrosine-protein kinase TEC is associated with bleeding due to its role in platelet aggregation^31, 32^.

**Figure 1.**
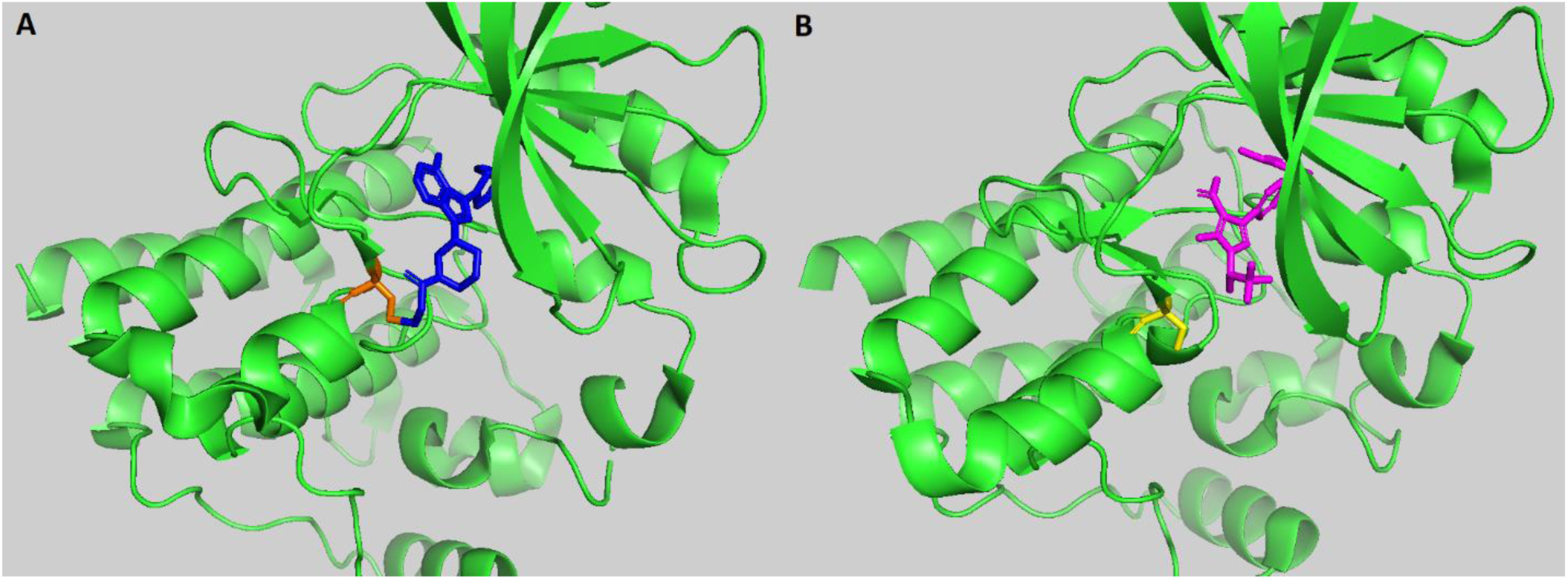
BTK inhibitors with different modes of action bound to BTK. A. Ibrutinib, covalently bound to the ATP-binding pocket of wild-type BTK. BTK is shown in green. Cysteine 481 is highlighted in orange. Ibrutinib is shown in dark blue. **B.** Pirtobrutinib, non-covalently bound to the ATP-binding pocket of mutant BTK C481S. BTK C481S is shown in green. Serine 481 is highlighted in yellow and Pirtobrutinib is in magenta. These images were created using PyMOL from the Protein Databank (PDB) entries 5P9J^33^ and 8FLN^25^, respectively.

## Results

In this study, we present a dataset of 15 BTK inhibitors that are currently FDA-approved or in clinical development. We provide resolution with respect to biochemical potency, selectivity, and biological effects of various BTK inhibitors, including the differential evaluation of covalent and non-covalent inhibitors. These insights help determine clinical dosing and may provide mechanistic clues as to why some drugs do better in the clinic than others.

### Most next-generation BTK inhibitors are more selective than ibrutinib

We utilized the scanMAX Kinase Profiling Panel (Eurofins Discovery, San Diego, CA) to determine the *in vitro* ATP-independent selectivity of 15 BTK inhibitors, when tested at 1 µM, to 468 kinases/kinase domains (409 of which are wild-type sequences and 403 are from unique proteins). This panel covers approximately 75% of the human kinome. A hit was counted when at least 65% of the kinase had been competed off the control ligand (a percent of control (POC) value <35) at the given concentration^34^. As expected, all the inhibitors tested hit BTK quite strongly (POC of 0 to 0.65) (Figure 2; Supplemental table 1). The total number of unique hits per inhibitor varied from 2 to 98 (Table 1, Figure 3). Most off-target hits observed were within the Tyrosine Kinase (TK) family (Figure 2). Remibrutinib, a covalent inhibitor, showed the most specificity, having only one off-target hit; while nemtabrutinib, a non-covalent inhibitor, showed the least specificity with 97 off-target hits (Table 1, Figure 3). Angst and colleagues found the same scanMAX specificity profile for 1 µM remibrutinib^35^. We categorized the selectivity of these inhibitors as high (≤11 off-targets), moderate (12-50 off-targets), and low (>50 off-targets) (Table 1). These categories did not correlate specifically with the mode of action of these inhibitors (Table 1; Figure 3).

**Figure 2.**
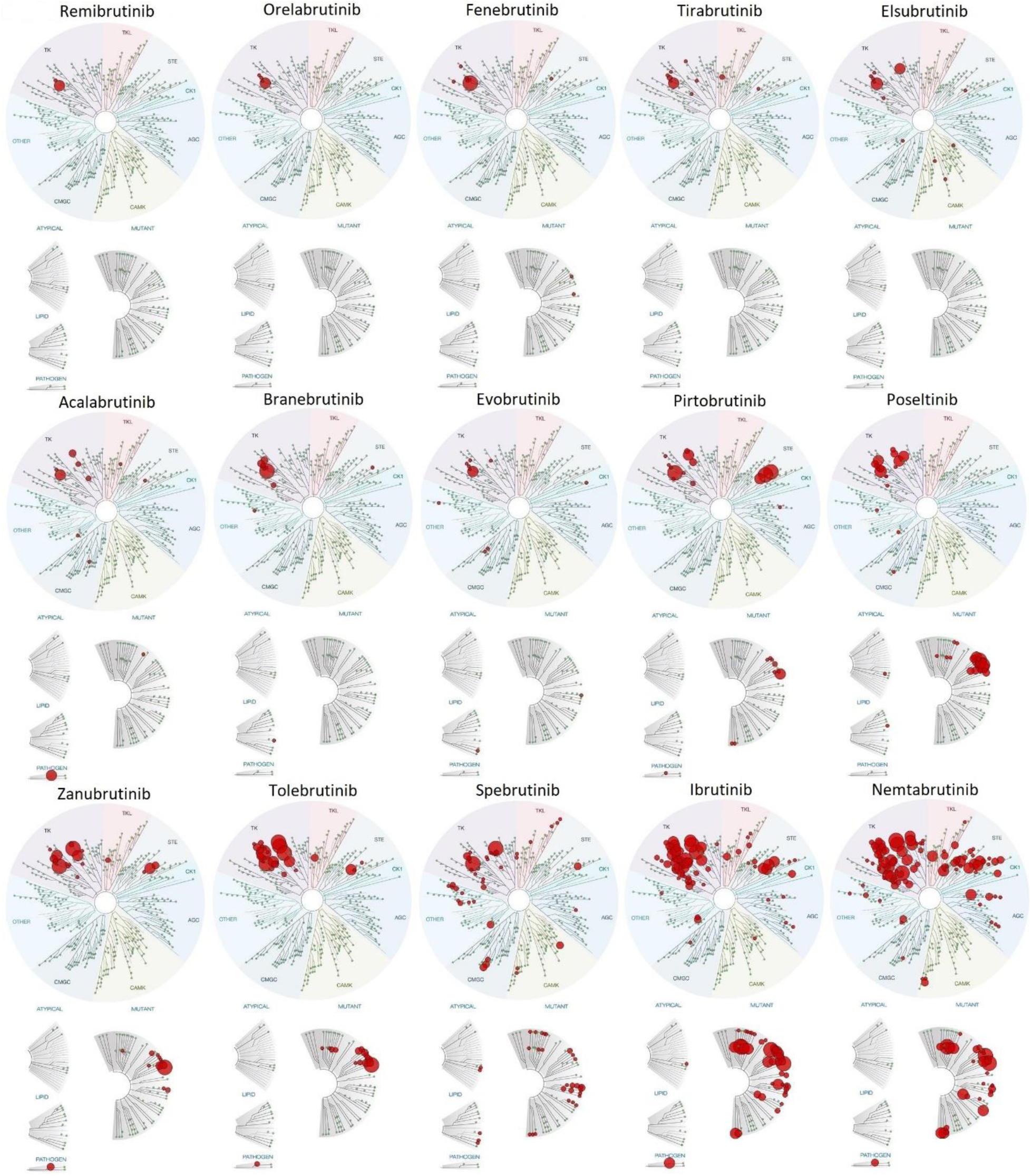
Kinome selectivity profile comparison of 15 known BTK inhibitors. TREEspot™ images showcasing the selectivity kinase panel hit profile for each BTK inhibitor as red dots on a phylogeny of kinases. The phylogenies are of wild-type, atypical, lipid, pathogen, and mutant kinases. Dot size indicates hit strength, so the larger the red dot, the stronger the hit. The BTK inhibitors are placed in order of most selective (top left) to least selective (bottom right).

**Figure 3.**
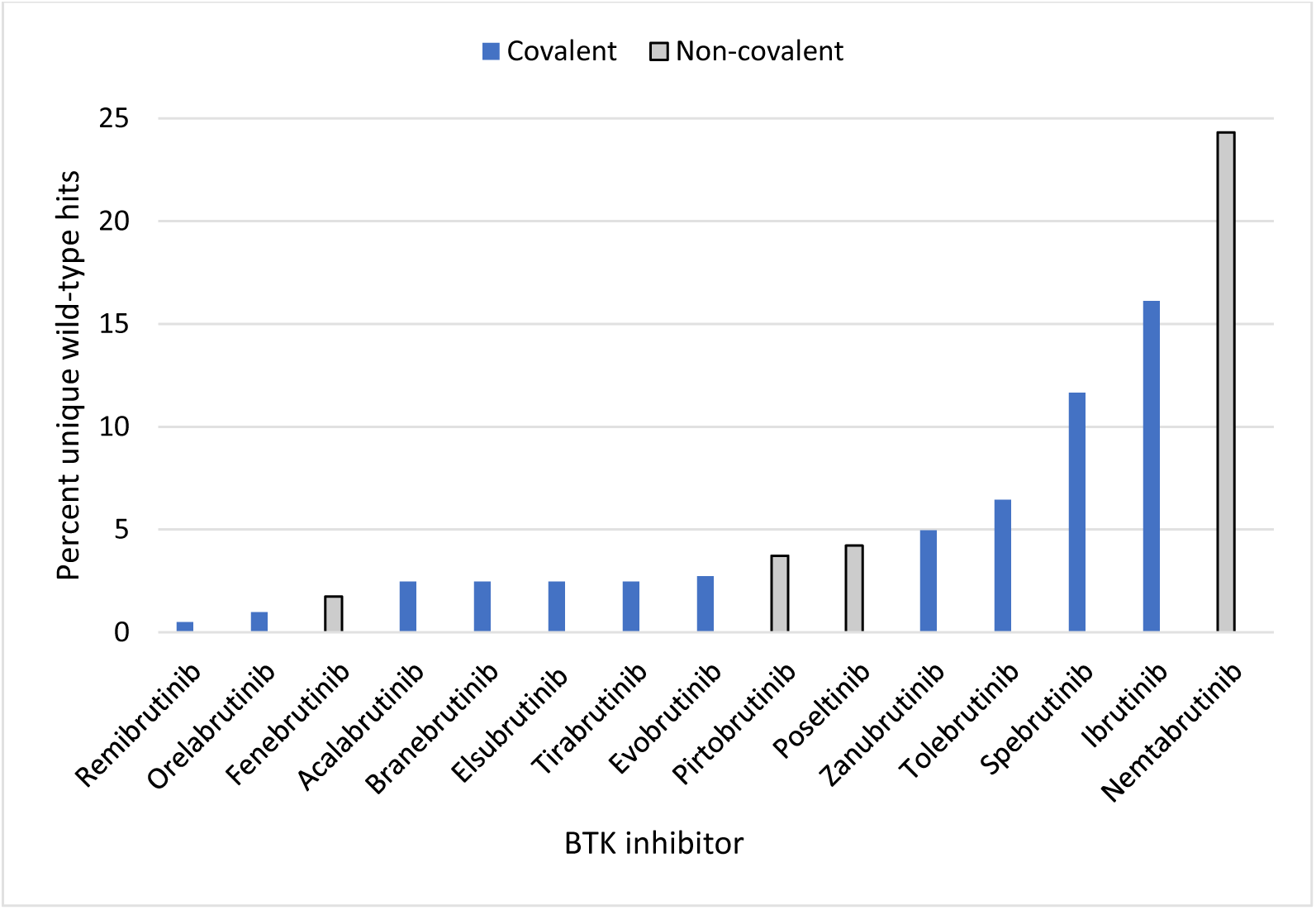
Selectivity ranking of included BTK inhibitors. The percentage of hits seen against the 403 unique wild- type kinases are plotted for each inhibitor tested. The compounds are plotted from the most specific (left) to least specific (right). Non-covalent compounds are highlighted (gray bars with black outline).

**Table 1.**
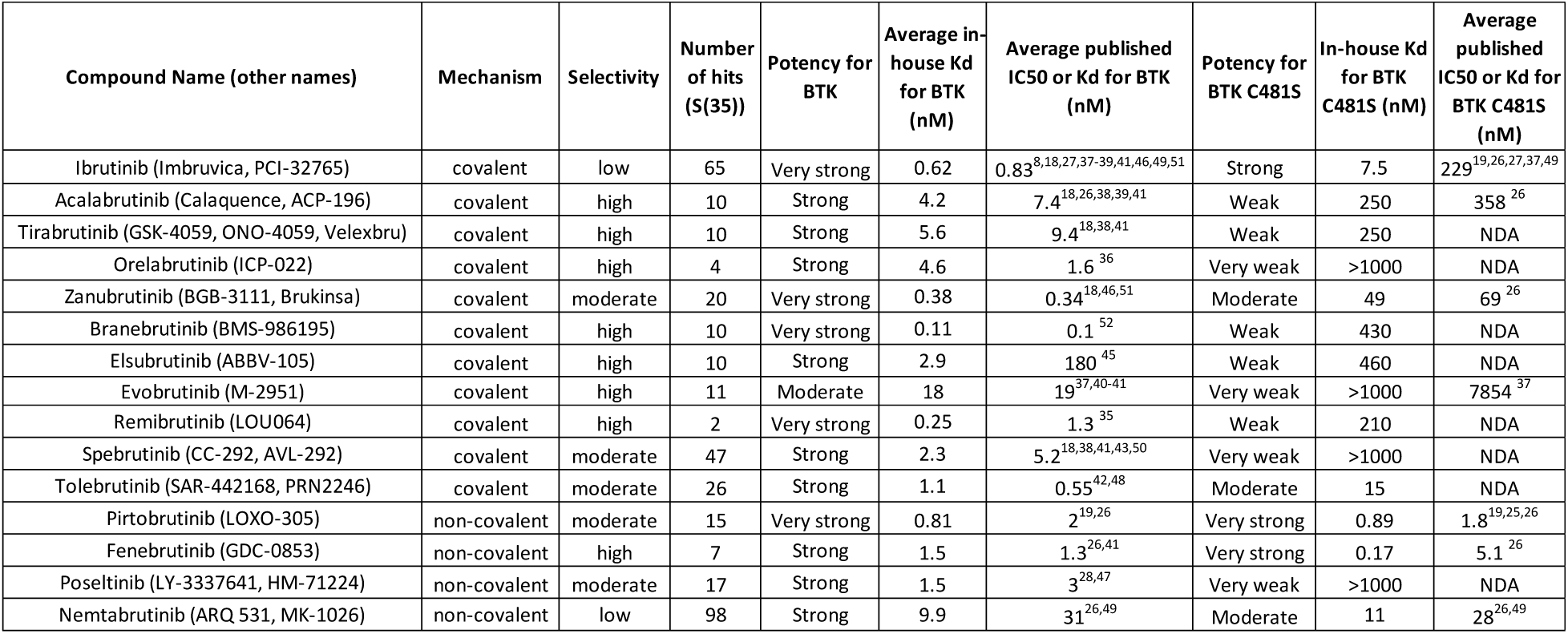
Summary of selectivity and binding affinity results for 15 known BTK inhibitors. The selectively score (S(35)) is the number of unique kinases (out of 403) that were competed off their control ligand by at least 65% at the screening concentration of 1 µM^34^. NDA: no data available. Published half-maximal inhibitory concentration (IC_50_) or dissociation constant (K_d_) values are denoted in the table^8, 18, 19, 25–28, 35–53^.

Our results correlate well with published results (e.g., ^18, 36, 37^). For example, ibrutinib bound to the four other TEC family kinases, interleukin-2-inducible T cell kinase (ITK/TSK), tyrosine kinase expressed in hepatocellular carcinoma (TEC), bone marrow-expressed kinase (BMX/ETK), and T cell-expressed kinase (TXK/RLK)^38^ (Supplemental table 1).

Most next-generation inhibitors bound to TEC and BMX with a similar POC as ibrutinib, but did not bind significantly to ITK (e.g., ^38, 54^) (Supplemental table 1). Additionally, the binding of TXK by next-generation BTK inhibitors was reduced when compared with that of ibrutinib (Supplemental table 1).

Moreover, ibrutinib bound to the other kinases with a cysteine in their active site, whereas most next-generation inhibitors had less specificity for many of these kinases (Supplemental table 2). This was especially so for mitogen-activated protein kinase kinase 7 (MKK7/SEK2/JNKK2), which was only found to be a weak hit for tolebrutinib and elsubrutinib (Supplemental table 2).

### No apparent correlation found between the potency of BTK inhibitors to wild-type BTK and their mode of action

Using Eurofins Discovery’s KdELECT Kinase Assay Panel (San Diego, CA), we measured the ATP-independent *in vitro* binding affinity of the aforementioned BTK inhibitors to wild-type BTK. Most of these BTK inhibitors bound to BTK with single digit nanomolar affinities (Figure 4; Supplemental table 3). The range of affinities for BTK was 0.11 nM to 18 nM (Figure 4; Supplemental table 3). Branebrutinib showed the strongest affinity and evobrutinib showed the weakest affinity (Figure 4). We categorized the affinity of each inhibitor into three groups: very strong (<1 nM), strong (1 nM to 10 nM), and moderate (10 nM to 100 nM) (Table 1). No apparent correlation between BTK binding affinity and BTK inhibitor binding mechanism was observed (Figure 4; Supplemental table 3).

**Figure 4.**
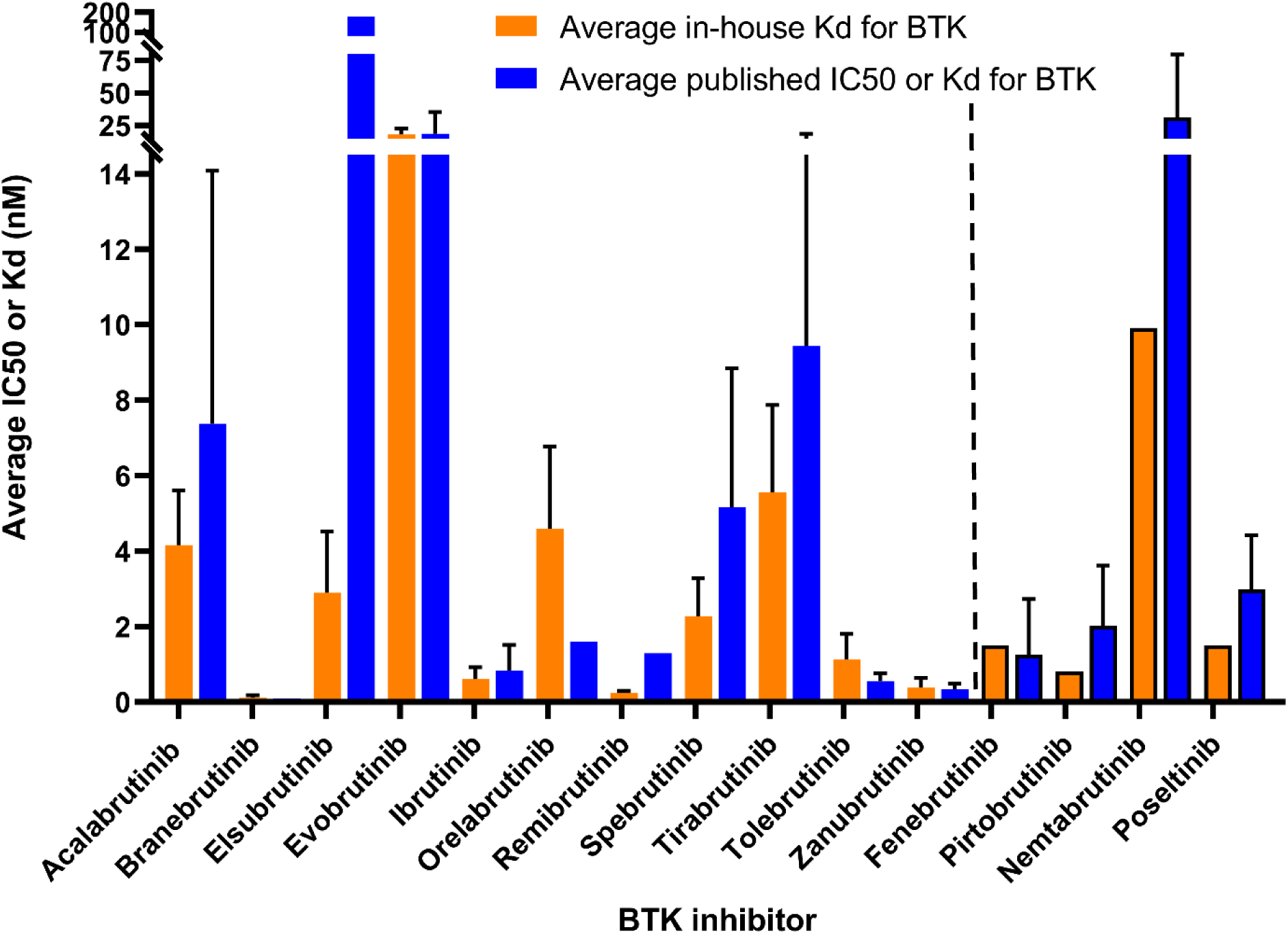
Comparison of averaged in-house K_d_ values with published IC_50_ or K_d_ values for BTK inhibitor binding to BTK. The first (orange) bar for each compound shows the in-house averaged K_d_ value (number of replicates is variable (Supplemental Table 3)). The second (blue) bar for each compound is the averaged published IC_50_ or K_d_ value(s) for each compound (number of replicates is variable (Supplemental Table 3)). Standard deviation error bars for those values generated from with more than one independent experiment are shown. Non-covalent compound results are highlighted to the right of the dashed vertical line with black outlines.

Most of our measured K_d_ values were within 3-fold of published IC_50_ or K_d_ values for these interactions (Figure 4; Supplemental table 3). The two exceptions were elsubrutinib and remibrutinib. The major discrepancy was for elsubrutinib where the only published IC_50_ value is 62 times weaker than the K_d_ value established from seven replicates^45^ (Figure 4; Supplemental table 3). This discrepancy may be due to differences in experimental design and/or detection limits of the assay. Goess and colleagues used an activity assay, whereas the KINOMEscan technology uses a binding competition assay. The single published IC_50_ value for remibrutinib is within 5.2-fold of the K_d_ value that we measured from seven replicates^35^ (Figure 4).

### Most non-covalent BTK inhibitors more potently bound the clinically relevant mutant BTK C481S protein than covalent BTK inhibitors

We measured the ATP-independent *in vitro* binding affinity of these BTK inhibitors to a clinically relevant resistance mutation of BTK (BTK C481S). The range of affinities for BTK C481S was 0.17 nM to >1 µM (Table 2). Fenebrutinib (GDC-0853), a non-covalent inhibitor, had the strongest affinity while evobrutinib, orelabrutinib, poseltinib, and spebrutinib (all covalent except poseltinib) did not demonstrate any binding at the concentrations tested (Table 2). We categorized the affinity of each inhibitor into five groups: very strong (<1 nM), strong (1 nM to 10 nM), moderate (10 nM to 100 nM), weak (100 nM to 1 µM), and very weak (>1 µM) (Table 1).

**Table 2.**
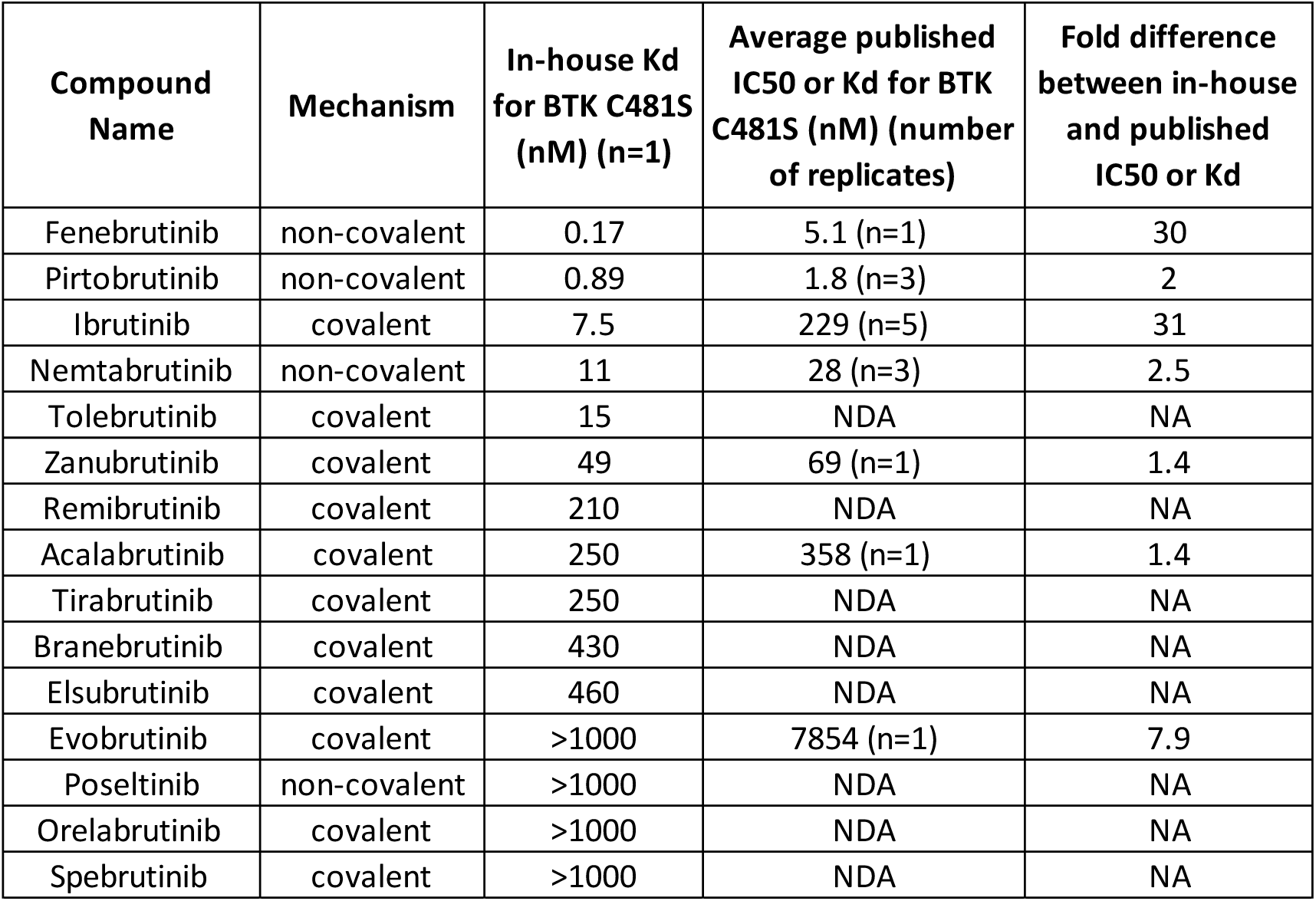
Comparison of in-house and published BTK inhibitor binding affinities for mutant BTK C481S. The mechanism of binding to wild-type BTK is listed in the second column. The BTK inhibitors are rank-ordered by our measured K_d_ values for BTK C481S. The number of replicates for BTK IC_50_ or K_d_ values is shown in parentheses after the IC_50_ or K_d_ value indicated. The fold-difference between measured and published results is shown in the fifth column. NDA: no data available. NA: not applicable.

The majority of non-covalent inhibitors tested showed stronger affinities for BTK C481S than covalent inhibitors (Table 2). However, three covalent inhibitors, ibrutinib, tolebrutinib, and zanubrutinib all had K_d_ values for BTK C481S of <50 nM (Table 2). We did not find many published IC_50_ or K_d_ values for this interaction but for those that are available, the majority were within 3-fold (Table 2).

The two exceptions to this were a 30-fold difference for fenebrutinib as compared with its published K_d_ value^26^ and a 31-fold difference for ibrutinib when compared with the average published binding measurements for it^19, 26, 27, 37, 49^ (Table 2).

We ranked the potency of these BTK inhibitors to wild-type BTK and BTK C481S and found that overall GDC-0853 (fenebrutinib), a non-covalent inhibitor, showed the strongest combined affinity to both proteins (Supplemental figure 1). Orelabrutinib, a covalent inhibitor, had the weakest total affinity for both proteins (Supplemental figure 1). Most of the non-covalent inhibitors evaluated had overall stronger combined affinities for these two forms of BTK than the covalent inhibitors tested (Supplemental figure 1). This is consistent with previous published results (e.g., ^16, 25, 49^).

### Non-covalent BTK inhibitors displayed fewer non-specific biological effects than covalent inhibitors

Testing the biological effects of a compound provides a valuable insight into the mechanism of action, most efficacious dose, and possible adverse effects that might be expected clinically^55–57^. We compared the biological effects elicited by ibrutinib, spebrutinib, pirtobrutinib, and fenebrutinib using the BioMAP® Diversity PLUS® Panel (Eurofins Discovery, St. Charles, MO). This panel includes 148 biomarkers in 12 human primary cell systems that model different aspects of human tissue biology and disease. No cytotoxic effects were observed for any of these compounds at the concentrations tested (Figure 5, no black arrows). All four BTK inhibitors showed antiproliferative activity in the T cell- activated B cell model (BT) at all four concentrations tested (Figure 5, gray arrows); likely through inhibition of BTK and thus the BCR signaling pathway. This is consistent with previous results (e.g., ^5, 25, 50, 58^). Moreover, all compounds demonstrated a decreased secretion of the survival signals interleukin- 6 (IL-6), interleukin-2 (IL-2), and tumor necrosis factor alpha (sTNF- α) in the B cell model of autoimmune disease, oncology, and inflammation (Figure 5).

**Figure 5.**
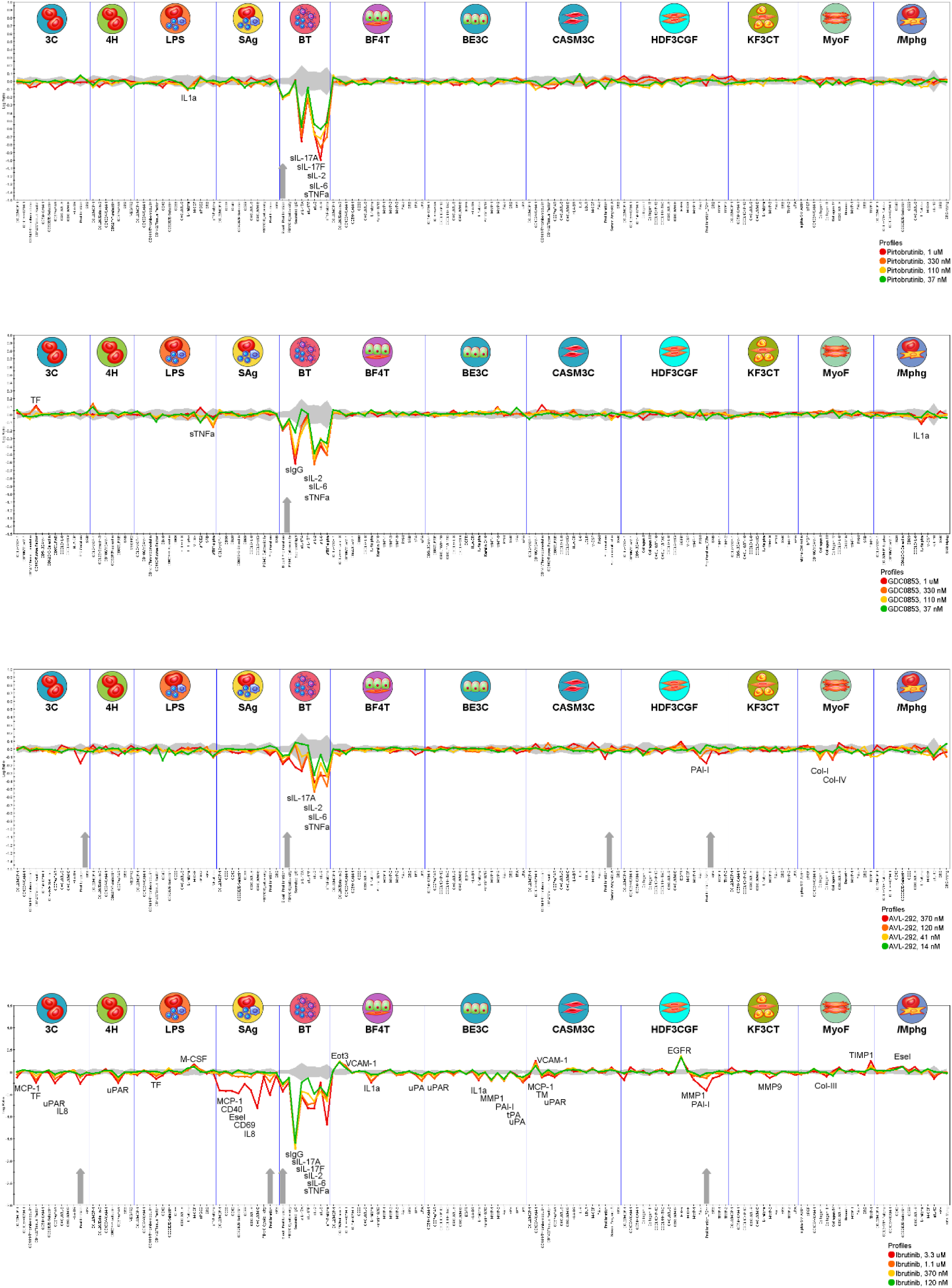
Biological effects of four BTK inhibitors in primary human cell-based models. Plots demonstrating the BioMAP® Diversity PLUS® Panel profile of treatment with pirtobrutinib, fenebrutinib (GDC0853), spebrutinib (AVL- 292), and ibrutinib. Each compound was tested at four concentrations (indicated by red, orange, yellow, and green traces). The 12 BioMAP systems are separated by vertical lines and contain representative icons at the top. Each biomarker is listed on the X-axis. Annotated peaks indicate statistically significant changes in biomarker readouts when compared with the vehicle control. Changes are displayed on the Y-axis as a log ratio when compared with the relevant vehicle control. The 95% confidence interval for the vehicle control is shown as a shaded gray area along the zero-change midline. Gray vertical arrows on the X-axis signify antiproliferative impact of the test drug at one or more concentrations; black vertical arrows (not present) would represent cytotoxic effects seen at one or more concentrations (i.e., none were seen in this study).

Overall, the two non-covalent inhibitors that we tested had the fewest biological effects, in terms of biomarkers modulated, and those effects were the most B cell model-specific (Figure 5). Pirtobrutinib had one less effect than fenebrutinib (GDC0853) and the fewest non-B cell effects (1 of 7 effects was not in the BT model versus 3 of 8 effects were not in the B cell model, respectively, when the inhibitors were tested at 1 µM) (Figure 5). The first-in-class BTK inhibitor, ibrutinib, showed the most biological effects (39 at 3.3 µM and 26 at 1.1 µM) and the most non-B cell model effects (32 at 3.3 µM and 19 at 1.1 µM), followed by spebrutinib (AVL-292) (11 at 370 nM, 6 of which were not in the BT model) (Figure 5).

While only minor differences were seen in inflammatory cytokine production and immune function between the non-covalent inhibitors and the second-generation covalent inhibitor (spebrutinib), a major difference observed was that treatment with spebrutinib decreased the activity of tissue remodeling biomarkers and produced antiproliferative effects in endothelial (3C), coronary artery smooth muscle (CASM), and fibroblast cells (HDF3CGF) (Figure 5, Table 3). These BioMAP results mostly correlate with the selectivity data generated (Figure 2). The exception is that we found more off-targets kinases for pirtobrutinib than for fenebrutinib (Table 1), including some of the kinases with a similar cysteine residue in their ATP-binding pockets (Supplemental tables 1 and 2).

**Table 3.**
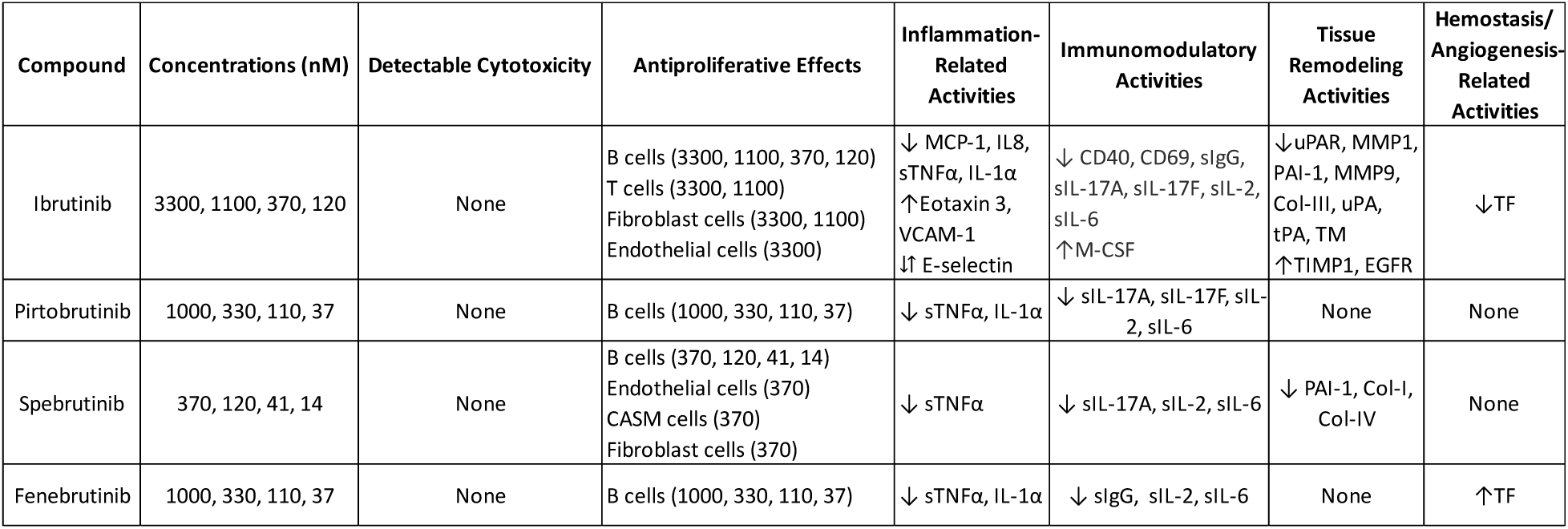
Summary of significant biological effects of ibrutinib, pirtobrutinib, spebrutinib, and fenebrutinib. ↑ signifies an increase in biomarker level relative to the vehicle control, while ↓indicates a decrease in biomarker level relative to the vehicle control, and ⇵ is used to show when an increase and a decrease was seen relative to vehicle control that occurred in different models.

Most next-generation inhibitor activities overlapped with that of ibrutinib (Figure 5; Table 3) and previous work^25, 41, 50^. A notable exception was the increase in the production of Tissue Factor (TF) in the 3C model (representing cardiovascular disease and chronic inflammation) following fenebrutinib treatment versus the decreased TF level seen after ibrutinib treatment in the same model (Figure 5; Table 3). Additionally, spebrutinib showed antiproliferative activity in CASM cells (modeling cardiovascular inflammation and restenosis), which was not seen with any other BTK inhibitor tested. This antiproliferative activity was not accompanied by any significant biomarker changes in this model (Figure 5). Spebrutinib also decreased the levels of different tissue remodeling biomarkers when compared with ibrutinib (i.e., collagen-I and IV versus collagen-III, respectively) (Figure 5; Table 3).

## Discussion

Accurately dissecting why drugs with the same target perform differently in clinical trials is challenging. Comparing drugs under the same conditions is a good way to tease out differences and gain a better understanding of each drug’s mechanism of action, biological outcomes, and potential adverse effects. Moreover, integration of specificity, potency, and biological effects early in drug discovery helps to quickly optimize therapeutic candidates. Here we measured the kinome-wide specificity and K_d_ values of 15 BTK inhibitors. We found a 96-fold difference in binding specificity, a 164-fold difference in wild-type BTK binding affinity, and a 2706-fold difference in the binding affinity for the clinically relevant mutant BTK C481S protein (Table 1). There was not a strong correlation between the binding mechanism and specificity (Figure 3) nor binding to wild-type BTK (Figure 4), however, non-covalent inhibitors generally bound more tightly to BTK C481S than covalent inhibitors (Table 2). This is consistent with clinical results, which show that non-covalent inhibitors can overcome some covalent inhibitor resistance (e.g., ^16, 17, 19, 49^). Moreover, we found differences in biological effects of three next-generation inhibitors, where the non-covalent inhibitors, pirtobrutinib and fenebrutinib, mainly showed only inflammation-related and immune-related effects, whereas the covalent inhibitor, spebrutinib, also showed tissue remodeling activity (Figure 5; Table 3). The different phenotypic profiles of covalent and non-covalent BTK inhibitors are likely due to secondary pharmacology effects, since they all have similar B cell-specific phenotypes that are likely due to the inhibition of BTK.

Most next-generation BTK inhibitors have successfully reduced the number of off-target interactions when compared with ibrutinib (Figure 2). Second-generation covalent inhibitors even have less specificity to the other TEC family members and other kinases with a cysteine residue in their ATP-binding site (Supplemental tables 1 and 2), consistent with published results (e.g., ^18^). This increased specificity is not accompanied by a loss of effectiveness of targeting BTK and treating B cell malignancies (e.g., ^59, 60^), which is consistent with the BioMAP human cellular phenotypic data generated in this study (Figure 5). Moreover, this increased specificity potentially makes these compounds better candidates for the treatment of autoimmune diseases^13^.

Interestingly, some of ibrutinib’s off-targets can be beneficial for the treatment of other diseases^22^. For example, the ability of ibrutinib to inhibit epidermal growth factor receptor (EGFR), human epidermal growth factor receptor 2 (HER2), Erb-B2 receptor tyrosine kinase 3 (ERBB3), and Erb- B2 receptor tyrosine kinase 4 (ERBB4) can be used to kill HER2+ breast cancer cells at a lower concentration than the established treatment with lapatinib^61^. This is consistent with our selectivity results, which showed that ibrutinib bound to EGFR, ERBB3, and ERBB4 to a similar extent as it did to BTK *in vitro* (Supplemental table 2). Wang and colleagues also found that spebrutinib (AVL-292) did not inhibit EGFR, HER2, ERBB3, and ERBB4, which is mostly consistent with our selectivity results. The one exception to this is that we found a moderate interaction (POC of 17) between spebrutinib and ERBB4 when tested at 1 µM (Supplemental table 2). Our EGFR and JAK3 specificity results are also consistent with *in vitro* and cellular data generated for spebrutinib and acalabrutinib^18, 43^.

The inhibition of multiple kinases, much like combination therapy using highly selective targeted drugs, was the idea behind the development of the non-covalent inhibitor nemtabrutinib^49^. Consistent with this, nemtabrutinib showed the most off-target interactions in our selectivity results (Figure 2; Table 1). Preclinical studies of nemtabrutinib showed promising results in mice models^49^. Reiff and colleagues identified an increase in survival in the Eμ-TCL1 engraftment model of CLL and an Eμ- MYC/TCL1 engraftment model resembling Richter’s transformation for nemtabrutinib versus ibrutinib-treated mice. It will be interesting to compare the effectiveness of nemtabrutinib with dual BTK inhibitors such as QL-X-138, a BTK/MNK (Mitogen-Activated Protein Kinase Interacting Kinase)^62^, in clinical trials.

We found that the strongest biological effects produced by the four BTK inhibitors tested were on B cells (Figure 5). This is expected when considering the function of BTK in B cells and published results (e.g., ^5, 25, 50, 58^). Additionally, spebrutinib’s effect on the level of sIL-17A produced, but not that of sIL-17F, is consistent with findings that these cytokines are regulated independently in B cells^63^ (Figure 5). A surprising finding for ibrutinib treatment was the increased production of EGFR in the HDF3CGF model (representing wound healing and inflammation) (Figure 5). Prior evidence (e.g., ^64, 65^) and our selectivity results (Supplemental table 2) support direct inhibition of EGFR by ibrutinib, thus a decrease in activity would be expected. However, ibrutinib treatment was also antiproliferative in this HDF3CGF model and decreased levels of the tissue remodelers MMP1 and PAI-I suggested that this increase in EGFR was not causing significant proliferative effects. It may also be that the increase in EGFR seen here represented the activation of a feedback loop in the cells to try and compensate for EGFR inhibition by the BTK inhibitor treatment. Another interesting finding is that all four BTK inhibitors tested decreased the production of TNF- α, a target for rheumatoid arthritis (RA), in the B cell model (Figure 5). This is not surprising since BTK is known to regulate TNF- α in RA (e.g., ^66^). What was unexpected is that fenebrutinib, but not spebrutinib, has shown significant efficacy in treating RA patients^50, 67^. Moreover, both ibrutinib and fenebrutinib block platelet aggregation from Fc-receptor CD32a (FcγRIIA) activation, which occurs in heparin-induced thrombocytopenia type II (HIT)^68^. Activated tissue factor (TF) expression is involved in the development of this condition; therefore, it is somewhat surprising that we found opposing effects of ibrutinib and fenebrutinib on TF activity in the 3C Th1 vasculature BioMAP system (Figure 5; Table 3). This suggests that these BTK inhibitors may have slightly different effects with respect to this condition, possibly through off-target effects or due to their different modes of action. Another consideration is that the effects of ibrutinib and fenebrutinib on TF levels was only seen at the higher concentrations of these inhibitors tested, whereas Goldmann and colleagues found that a low concentration of these inhibitors is sufficient for blocking platelet aggregation and thus making them potential candidates for HIT treatment.

Even though BTK inhibitors were the primary focus of this study, the screening cascade and various technologies utilized here may be applied to other target classes and chemical matter in a similar fashion. These preclinical tools thus provide the ability to conduct in-depth head-to-head comparisons of drug candidates, yielding valuable insights into a drug’s engagement with its target of interest and off-targets that could affect a broad range of biological effects, which may ultimately affect both efficacy and safety in the clinical therapeutic setting.

## Methods

### Small Molecules

Ibrutinib, Acalabrutinib, Tirabrutinib, Orelabrutinib, Zanubrutinib, Pirtobrutinib, Nemtabrutinib, Fenebrutinib, Poseltinib, Spebrutinib, Branebrutinib, Evobrutinib, Elsubrutinib, Tolebrutinib, and Remibrutinib were purchased from MedChemExpress (Monmouth Junction, NJ).

### Protein Constructs and Protein Expression

Full length wild-type BTK and mutant BTK C481S constructs were fused N-terminally with the DNA binding domain of NFκB (consisting of residues 35-36 fused to residues 41-359 (as described in ^69^), using UniProt entry P19838 as a reference) and expressed in transiently transfected HEK293 cells. Protein extracts were harvested in M-PER extraction buffer (Pierce Biotechnology, Rockford, IL) supplemented with 150 mM NaCl, 10 mM Dithithreitol (DTT), Protease Inhibitor Cocktail Complete (Roche Diagnostics GmbH, Mannheim, Germany) and Phosphatase Inhibitor Cocktail Set II (Merck KGaA, Darmstadt, Germany) following the manufacturer’s guidelines. The fusion protein was labeled with a DNA tag containing the NFκB binding site fused to an amplicon for qPCR readout, which was added directly to the expression extract.

### Competition Binding Assays

Small molecule binding to wild-type and mutant kinases was assessed using ATP site-dependent competition binding assays as previously described^34, 70–72^. Streptavidin-coated magnetic beads (Thermo Fisher Scientific, Waltham, MA) were incubated with a biotinylated affinity ligand for 30 min at 25°C. Liganded beads were blocked with excess biotin (125 nM) and washed with a blocking buffer containing SeaBlock (Pierce Biotechnology), 1% Bovine Serum Albumin (BSA) and 0.05% Tween 20. The binding reactions were prepared with the DNA tagged BTK protein extract, ligated affinity beads, and the competitor test compounds in a binding buffer (1x Phosphate Buffered Saline (PBS), 0.05% Tween 20, 10 mM DTT, 0.1% BSA, 2 mg/ml sheared salmon sperm DNA) in deep well, natural polypropylene 384-well plates, catalog number 784201 (Greiner Bio-One, Kremsmünster, Austria) in a final volume of 19.7 µL. These test compounds were prepared as 111x stocks in 100% Dimethyl sulfoxide (DMSO). All compounds for K_d_ measurements are distributed by acoustic transfer (non-contact dispensing) in 100% DMSO. The compounds were diluted directly into the assays such that the final concentration of DMSO is 0.9%. No enzyme purification steps were performed on the protein extracts before adding them to the reaction mixture, and the protein extracts were diluted 10,000-fold in the final reaction mixture (the final DNA-tagged enzyme concentration was less than 0.1 nM). Binding assay mixtures were incubated at 25 °C with shaking for 1 hr. Then affinity beads were extensively treated with a wash buffer (1x PBS, 0.05% Tween 20) to remove unbound protein from the protein lysate. Using an elution buffer (1x PBS, 0.05% Tween 20 and either 0.5 μM affinity ligand, the beads were resuspended and incubated at 25 °C while shaking for a 30 min period. The concentration of wild-type or mutant BTK in the eluates was then determined by quantitative PCR. K_d_ values for each competitor compound were determined using eleven serial threefold dilutions and three DMSO control points. Each assays affinity ligand concentration on the magnetic beads has been optimized to ensure that the true thermodynamic K_d_ values for competitor molecules were measured, as described in detail previously^72^.

### Data Analysis for Competitive Binding Assays

K_d_s were calculated using a standard dose-response curve fitting of the data using the Hill equation: Response = Background + (Signal – Background) / (1 + (K_dHill Slope_ / Dose^Hill^ ^Slope^). The Hill Slope was set to -1. A non-linear least square fit using the Levenberg-Marquardt algorithm was employed for curve fitting.

### Human primary cell systems

BioMAP® Diversity PLUS® Panel (Eurofins Discovery, St. Charles, MO) is composed of 12 human primary cell cultures or cocultures. The systems and stimuli have been described previously^73^. Adherent cells were added to 96-well plates, allowed to reach confluence, and incubated for 1 hr with the indicated BTK inhibitor or assay control, all dissolved in 0.1% DMSO. The cells were then stimulated with cytokines, agonists, or growth factors and cultured for a minimum of 24 hrs at 37°C in 5% CO_2_. Assay controls included DMSO-only (vehicle control), non-stimulated conditions (negative control), and a positive control test agent (colchicine at 1μM).

### Measured biomarkers

Following the system-specific incubation time, a total of 148 protein biomarkers spanning all 12 cell systems were measured, either in cellular extracts by direct ELISA, or in supernatants by bead-based multiplex immunoassays, homogeneous time-resolved fluorescence (HTRF) detection or capture ELISA. Cell proliferation and viability were measured using 0.1% sulforhodamine B (SRB, Sigma– Aldrich) after fixation with 10% TCA, and reading wells at 560 nm for adherent cells, and alamarBlue (Bio-Rad) for suspension cells. These biomarkers were previously described by Shah and colleagues.

### Biomarker analysis

For each biomarker, the value obtained from the BTK inhibitor-treated samples was divided by the average of 8 DMSO-only controls from the same plate, yielding a ratio that was then transformed to the log_10_ scale. Data from historical controls of prior BioMAP studies were used to define a range (95% confidence interval) for each ratio called “envelope.” Experimental ratios that decreased more than 20% (<0.097 on the log_10_ scale) or increased more than 20% (>0.079 on the log_10_ scale) when compared with the envelope were considered significant, and annotated when observed for 2 or more consecutive concentrations. For cytotoxicity, we considered a significant decrease in the SRB readout greater than 50% (<−0.3 on the log_10_ scale) and excluded from the analysis the corresponding concentrations. The acceptance criteria and quality assurance of the BioMAP experiments have been previously described^73^.

## Acknowledgements

The authors acknowledge members of Eurofins Discovery for their help with initial discussions surrounding the project and for feedback on the manuscript. The Table of Contents graphic was generated using BioRender.com. The ibrutinib structure in the Table of Contents graphic was obtained from PubChem (https://pubchem.ncbi.nlm.nih.gov/compound/24821094#section=2D-Structure). The pirtobrutinib structure in the Table of Contents graphic was obtained from PubChem (https://pubchem.ncbi.nlm.nih.gov/compound/129269915#section=2D-Structure).

## Author contributions

A.C.D. analyzed the specificity and dissociation constant data, contributed to the BioMAP analysis, and wrote the manuscript. A.M.H. contributed to the generation of the specificity and dissociation constant data. M.J. and N.B.S. contributed to the developed of the specificity and dissociation constant experiments. J.H.L. contributed to the generation and analysis of the BioMAP data. A.J.K. contributed to the development of the BioMAP experiments. All authors reviewed the manuscript.

## Supporting Information

Supplemental Table 1. Comparison of BTK inhibitor specificity to TEC family members. Supplemental Table 3. Comparison of in-house and published BTK inhibitor binding affinity to BTK. Supplemental Figure 1. Potency curves of 15 BTK inhibitors to BTK and mutant BTK (BTK C481S).

## References

[1] Cohen, P., Cross, D., and Jänne, P. A. (2021) Kinase drug discovery 20 years after imatinib: progress and future directions, Nat Rev Drug Discov 20, 551–569.

[2] Zarrin, A. A., Bao, K., Lupardus, P., and Vucic, D. (2021) Kinase inhibition in autoimmunity and inflammation, Nat Rev Drug Discov 20, 39–63.

[3] Smith, C. I., Islam, T. C., Mattsson, P. T., Mohamed, A. J., Nore, B. F., Vihinen, M., Shah, F., Stepan, A. F., O’Mahony, A., Velichko, S., Folias, A. E., Houle, C., Shaffer, C. L., Marcek, J., Whritenour, J., Stanton, R., Berg, E. L., Schafer, P. H., Kivitz, A. J., Ma, J., Korish, S., Sutherland, D., Li, L., Azaryan, A., Kosek, J., Adams, M., Capone, L., Hur, E. M., Hough, D. R., and Ringheim, G. E. (2001) The Tec family of cytoplasmic tyrosine kinases: mammalian Btk, Bmx, Itk, Tec, Txk and homologs in other species Mechanisms of Skin Toxicity Associated with Metabotropic Glutamate Receptor 5 Negative Allosteric Modulators Spebrutinib (CC-292) Affects Markers of B Cell Activation, Chemotaxis, and Osteoclasts in Patients with Rheumatoid Arthritis: Results from a Mechanistic Study, Bioessays 23, 436-446.

4. [4] Hendriks, R. W., de Bruijn, M. F., Maas, A., Dingjan, G. M., Karis, A., and Grosveld, F. (1996) Inactivation of Btk by insertion of lacZ reveals defects in B cell development only past the pre-B cell stage, EMBO J 15, 4862–4872.

[5] Herman, S. E., Gordon, A. L., Hertlein, E., Ramanunni, A., Zhang, X., Jaglowski, S., Flynn, J., Jones, J., Blum, K. A., Buggy, J. J., Hamdy, A., Johnson, A. J., and Byrd, J. C. (2011) Bruton tyrosine kinase represents a promising therapeutic target for treatment of chronic lymphocytic leukemia and is effectively targeted by PCI-32765, Blood 117, 6287–6296.

[6] Pal Singh, S., Dammeijer, F., and Hendriks, R. W. (2018) Role of Bruton’s tyrosine kinase in B cells and malignancies, Mol Cancer 17, 57.

[7] Woyach, J. A., Bojnik, E., Ruppert, A. S., Stefanovski, M. R., Goettl, V. M., Smucker, K. A., Smith, L. L., Dubovsky, J. A., Towns, W. H., MacMurray, J., Harrington, B. K., Davis, M. E., Gobessi, S., Laurenti, L., Chang, B. Y., Buggy, J. J., Efremov, D. G., Byrd, J. C., and Johnson, A. J. (2014) Bruton’s tyrosine kinase (BTK) function is important to the development and expansion of chronic lymphocytic leukemia (CLL), Blood 123, 1207–1213.

[8] Honigberg, L. A., Smith, A. M., Sirisawad, M., Verner, E., Loury, D., Chang, B., Li, S., Pan, Z., Thamm, D. H., Miller, R. A., and Buggy, J. J. (2010) The Bruton tyrosine kinase inhibitor PCI-32765 blocks B-cell activation and is efficacious in models of autoimmune disease and B-cell malignancy, Proc Natl Acad Sci U S A 107, 13075–13080.

[9] Broccoli, A., Argnani, L., Morigi, A., Nanni, L., Casadei, B., Pellegrini, C., Stefoni, V., and Zinzani, P. L. (2021) Long-Term Efficacy and Safety of Ibrutinib in the Treatment of CLL Patients: A Real Life Experience, J Clin Med 10.

[10] O’Brien, S., Furman, R. R., Coutre, S. E., Sharman, J. P., Burger, J. A., Blum, K. A., Grant, B., Richards, D. A., Coleman, M., Wierda, W. G., Jones, J. A., Zhao, W., Heerema, N. A., Johnson, A. J., Izumi, R., Hamdy, A., Chang, B. Y., Graef, T., Clow, F., Buggy, J. J., James, D. F., and Byrd, J. C. (2014) Ibrutinib as initial therapy for elderly patients with chronic lymphocytic leukaemia or small lymphocytic lymphoma: an open-label, multicentre, phase 1b/2 trial, Lancet Oncol 15, 48–58.

[11] Wang, M. L., Rule, S., Martin, P., Goy, A., Auer, R., Kahl, B. S., Jurczak, W., Advani, R. H., Romaguera, J. E., Williams, M. E., Barrientos, J. C., Chmielowska, E., Radford, J., Stilgenbauer, S., Dreyling, M., Jedrzejczak, W. W., Johnson, P., Spurgeon, S. E., Li, L., Zhang, L., Newberry, K., Ou, Z., Cheng, N., Fang, B., McGreivy, J., Clow, F., Buggy, J. J., Chang, B. Y., Beaupre, D. M., Kunkel, L. A., and Blum, K. A. (2013) Targeting BTK with ibrutinib in relapsed or refractory mantle-cell lymphoma, N Engl J Med 369, 507–516.

[12] Xiao, L., Salem, J. E., Clauss, S., Hanley, A., Bapat, A., Hulsmans, M., Iwamoto, Y., Wojtkiewicz, G., Cetinbas, M., Schloss, M. J., Tedeschi, J., Lebrun-Vignes, B., Lundby, A., Sadreyev, R. I., Moslehi, J., Nahrendorf, M., Ellinor, P. T., and Milan, D. J. (2020) Ibrutinib-Mediated Atrial Fibrillation Attributable to Inhibition of C-Terminal Src Kinase, Circulation 142, 2443–2455.

[13] Alu, A., Lei, H., Han, X., Wei, Y., and Wei, X. (2022) BTK inhibitors in the treatment of hematological malignancies and inflammatory diseases: mechanisms and clinical studies, J Hematol Oncol 15, 138.

[14] Ahn, I. E., Underbayev, C., Albitar, A., Herman, S. E., Tian, X., Maric, I., Arthur, D. C., Wake, L., Pittaluga, S., Yuan, C. M., Stetler-Stevenson, M., Soto, S., Valdez, J., Nierman, P., Lotter, J., Xi, L., Raffeld, M., Farooqui, M., Albitar, M., and Wiestner, A. (2017) Clonal evolution leading to ibrutinib resistance in chronic lymphocytic leukemia, Blood 129, 1469–1479.

[15] Maddocks, K. J., Ruppert, A. S., Lozanski, G., Heerema, N. A., Zhao, W., Abruzzo, L., Lozanski, A., Davis, M., Gordon, A., Smith, L. L., Mantel, R., Jones, J. A., Flynn, J. M., Jaglowski, S. M., Andritsos, L. A., Awan, F., Blum, K. A., Grever, M. R., Johnson, A. J., Byrd, J. C., and Woyach, J. A. (2015) Etiology of Ibrutinib Therapy Discontinuation and Outcomes in Patients With Chronic Lymphocytic Leukemia, JAMA Oncol 1, 80–87.

[16] Naeem, A., Utro, F., Wang, Q., Cha, J., Vihinen, M., Martindale, S., Zhou, Y., Ren, Y., Tyekucheva, S., Kim, A. S., Fernandes, S. M., Saksena, G., Rhrissorrakrai, K., Levovitz, C., Danysh, B. P., Slowik, K., Jacobs, R. A., Davids, M. S., Lederer, J. A., Zain, R., Smith, C. I. E., Leshchiner, I., Parida, L., Getz, G., and Brown, J. R. (2023) Pirtobrutinib targets BTK C481S in ibrutinib-resistant CLL but second-site BTK mutations lead to resistance, Blood Adv 7, 1929–1943.

[17] Quinquenel, A., Fornecker, L. M., Letestu, R., Ysebaert, L., Fleury, C., Lazarian, G., Dilhuydy, M. S., Nollet, D., Guieze, R., Feugier, P., Roos-Weil, D., Willems, L., Michallet, A. S., Delmer, A., Hormigos, K., Levy, V., Cymbalista, F., Baran-Marszak, F., Thompson, P. A., and Tam, C. S. (2019) Prevalence of BTK and PLCG2 mutations in a real-life CLL cohort still on ibrutinib after 3 years: a FILO group study Pirtobrutinib: a new hope for patients with BTK inhibitor-refractory lymphoproliferative disorders, Blood 134, 641–644.

18. Barf, T., Covey, T., Izumi, R., van de Kar, B., Gulrajani, M., van Lith, B., van Hoek, M., de Zwart, E., Mittag, D., Demont, D., Verkaik, S., Krantz, F., Pearson, P. G., Ulrich, R., and Kaptein, A. (2017) Acalabrutinib (ACP-196): A Covalent Bruton Tyrosine Kinase Inhibitor with a Differentiated Selectivity and In Vivo Potency Profile, J Pharmacol Exp Ther 363, 240-252.

[19] Brandhuber, B., Gomez, E., Smith, S., Eary, T., Spencer, S., Rothenberg, S. M., and Andrews, S. (2018) LOXO-305, a next generation reversible BTK inhibitor, for overcoming acquired resistance to irreversible BTK inhibitors, *Clinical Lymphoma*, Myeloma and Leukemia 18.

[20] Wen, T., Wang, J., Shi, Y., Qian, H., and Liu, P. (2021) Inhibitors targeting Bruton’s tyrosine kinase in cancers: drug development advances, Leukemia 35, 312–332.

[21] Dhillon, S., Wu, H., Hu, C., Wang, A., Weisberg, E. L., Chen, Y., Yun, C. H., Wang, W., Liu, Y., Liu, X., Tian, B., Wang, J., Zhao, Z., Liang, Y., Li, B., Wang, L., Wang, B., Chen, C., Buhrlage, S. J., Qi, Z., Zou, F., Nonami, A., Li, Y., Fernandes, S. M., Adamia, S., Stone, R. M., Galinsky, I. A., Wang, X., Yang, G., Griffin, J. D., Brown, J. R., Eck, M. J., Liu, J., Gray, N. S., and Liu, Q. (2020) Tirabrutinib: First Approval Discovery of a BTK/MNK dual inhibitor for lymphoma and leukemia, Drugs 80, 835–840.

[22] Rozkiewicz, D., Hermanowicz, J. M., Kwiatkowska, I., Krupa, A., and Pawlak, D. (2023) Bruton’s Tyrosine Kinase Inhibitors (BTKIs): Review of Preclinical Studies and Evaluation of Clinical Trials, Molecules 28.

[23] Dhillon, S., Dhillon, S., Wu, H., Hu, C., Wang, A., Weisberg, E. L., Chen, Y., Yun, C. H., Wang, W., Liu, Y., Liu, X., Tian, B., Wang, J., Zhao, Z., Liang, Y., Li, B., Wang, L., Wang, B., Chen, C., Buhrlage, S. J., Qi, Z., Zou, F., Nonami, A., Li, Y., Fernandes, S. M., Adamia, S., Stone, R. M., Galinsky, I. A., Wang, X., Yang, G., Griffin, J. D., Brown, J. R., Eck, M. J., Liu, J., Gray, N. S., and Liu, Q. (2021) Orelabrutinib: First Approval Tirabrutinib: First Approval Discovery of a BTK/MNK dual inhibitor for lymphoma and leukemia, Drugs 81, 503–507.

[24] Thompson, P. A., and Tam, C. S. (2023) Pirtobrutinib: a new hope for patients with BTK inhibitor-refractory lymphoproliferative disorders, Blood 141, 3137–3142.

[25] Gomez, E. B., Ebata, K., Randeria, H. S., Rosendahl, M. S., Cedervall, E. P., Morales, T. H., Hanson, L. M., Brown, N. E., Gong, X., Stephens, J., Wu, W., Lippincott, I., Ku, K. S., Walgren, R. A., Abada, P. B., Ballard, J. A., Allerston, C. K., and Brandhuber, B. J. (2023) Preclinical characterization of pirtobrutinib, a highly selective, noncovalent (reversible) BTK inhibitor, Blood 142, 62–72.

[26] Wang, E., Mi, X., Thompson, M. C., Montoya, S., Notti, R. Q., Afaghani, J., Durham, B. H., Penson, A., Witkowski, M. T., Lu, S. X., Bourcier, J., Hogg, S. J., Erickson, C., Cui, D., Cho, H., Singer, M., Totiger, T. M., Chaudhry, S., Geyer, M., Alencar, A., Linley, A. J., Palomba, M. L., Coombs, C. C., Park, J. H., Zelenetz, A., Roeker, L., Rosendahl, M., Tsai, D. E., Ebata, K., Brandhuber, B., Hyman, D. M., Aifantis, I., Mato, A., Taylor, J., and Abdel-Wahab, O. (2022) Mechanisms of Resistance to Noncovalent Bruton’s Tyrosine Kinase Inhibitors, N Engl J Med 386, 735–743.

[27] Furman, R. R., Cheng, S., Lu, P., Setty, M., Perez, A. R., Guo, A., Racchumi, J., Xu, G., Wu, H., Ma, J., Steggerda, S. M., Coleman, M., Leslie, C., and Wang, Y. L. (2014) Ibrutinib resistance in chronic lymphocytic leukemia, N Engl J Med 370, 2352–2354.

[28] Byun, J. Y., Koh, Y. T., Jang, S. Y., Witcher, J. W., Chan, J. R., Pustilnik, A., Daniels, M. J., Kim, Y. H., Suh, K. H., Linnik, M. D., and Lee, Y. M. (2021) Target modulation and pharmacokinetics/pharmacodynamics translation of the BTK inhibitor poseltinib for model-informed phase II dose selection, Sci Rep 11, 18671.

[29] Ringheim, G. E., Wampole, M., and Oberoi, K. (2021) Bruton’s Tyrosine Kinase (BTK) Inhibitors and Autoimmune Diseases: Making Sense of BTK Inhibitor Specificity Profiles and Recent Clinical Trial Successes and Failures, Front Immunol 12, 662223.

[30] Lipsky, A., and Lamanna, N. (2020) Managing toxicities of Bruton tyrosine kinase inhibitors, Hematology Am Soc Hematol Educ Program 2020, 336–345.

[31] Atkinson, B. T., Ellmeier, W., and Watson, S. P. (2003) Tec regulates platelet activation by GPVI in the absence of Btk, Blood 102, 3592–3599.

[32] Hamazaki, Y., Kojima, H., Mano, H., Nagata, Y., Todokoro, K., Abe, T., Nagasawa, T., Haselmayer, P., Camps, M., Liu-Bujalski, L., Nguyen, N., Morandi, F., Head, J., O’Mahony, A., Zimmerli, S. C., Bruns, L., Bender, A. T., Schroeder, P., and Grenningloh, R. (1998) Tec is involved in G protein-coupled receptor- and integrin-mediated signalings in human blood platelets Efficacy and Pharmacodynamic Modeling of the BTK Inhibitor Evobrutinib in Autoimmune Disease Models, Oncogene 16, 2773–2779.

[33] Bender, A. T., Gardberg, A., Pereira, A., Johnson, T., Wu, Y., Grenningloh, R., Head, J., Morandi, F., Haselmayer, P., and Liu-Bujalski, L. (2017) Ability of Bruton’s Tyrosine Kinase Inhibitors to Sequester Y551 and Prevent Phosphorylation Determines Potency for Inhibition of Fc Receptor but not B-Cell Receptor Signaling, Mol Pharmacol 91, 208–219.

[34] Karaman, M. W., Herrgard, S., Treiber, D. K., Gallant, P., Atteridge, C. E., Campbell, B. T., Chan, K. W., Ciceri, P., Davis, M. I., Edeen, P. T., Faraoni, R., Floyd, M., Hunt, J. P., Lockhart, D. J., Milanov, Z. V., Morrison, M. J., Pallares, G., Patel, H. K., Pritchard, S., Wodicka, L. M., Zarrinkar, P. P., Isenberg, D., Furie, R., Jones, N. S., Guibord, P., Galanter, J., Lee, C., McGregor, A., Toth, B., Rae, J., Hwang, O., Desai, R., Lokku, A., Ramamoorthi, N., Hackney, J. A., Miranda, P., de Souza, V. A., Jaller-Raad, J. J., Maura Fernandes, A., Garcia Salinas, R., Chinn, L. W., Townsend, M. J., Morimoto, A. M., Tuckwell, K., Jarmoskaite, I., AlSadhan, I., Vaidyanathan, P. P., and Herschlag, D. (2008) A quantitative analysis of kinase inhibitor selectivity Efficacy, Safety, and Pharmacodynamic Effects of the Bruton’s Tyrosine Kinase Inhibitor Fenebrutinib (GDC-0853) in Systemic Lupus Erythematosus: Results of a Phase II, Randomized, Double-Blind, Placebo-Controlled Trial How to measure and evaluate binding affinities, Nat Biotechnol 26, 127-132.

[35] Angst, D., Gessier, F., Janser, P., Vulpetti, A., Walchli, R., Beerli, C., Littlewood-Evans, A., Dawson, J., Nuesslein-Hildesheim, B., Wieczorek, G., Gutmann, S., Scheufler, C., Hinniger, A., Zimmerlin, A., Funhoff, E. G., Pulz, R., and Cenni, B. (2020) Discovery of LOU064 (Remibrutinib), a Potent and Highly Selective Covalent Inhibitor of Bruton’s Tyrosine Kinase, J Med Chem 63, 5102–5118.

[36] Cao, X. X., Jin, J., Fu, C. C., Yi, S. H., Zhao, W. L., Sun, Z. M., Yang, W., Li, D. J., Cui, G. H., Hu, J. D., Liu, T., Song, Y. P., Xu, B., Zhu, Z. M., Xu, W., Zhang, M. Z., Tian, Y. M., Zhang, B., Zhao, R. B., and Zhou, D. B. (2022) Evaluation of orelabrutinib monotherapy in patients with relapsed or refractory Waldenström’s macroglobulinemia in a single-arm, multicenter, open-label, phase 2 study, EClinicalMedicine 52, 101682.

[37] Haselmayer, P., Camps, M., Liu-Bujalski, L., Nguyen, N., Morandi, F., Head, J., O’Mahony, A., Zimmerli, S. C., Bruns, L., Bender, A. T., Schroeder, P., and Grenningloh, R. (2019) Efficacy and Pharmacodynamic Modeling of the BTK Inhibitor Evobrutinib in Autoimmune Disease Models, J Immunol 202, 2888–2906.

[38] Liclican, A., Serafini, L., Xing, W., Czerwieniec, G., Steiner, B., Wang, T., Brendza, K. M., Lutz, J. D., Keegan, K. S., Ray, A. S., Schultz, B. E., Sakowicz, R., Feng, J. Y., Kim, Y. Y., Park, K. T., Jang, S. Y., Lee, K. H., Byun, J. Y., Suh, K. H., Lee, Y. M., Kim, Y. H., and Hwang, K. W. (2020) Biochemical characterization of tirabrutinib and other irreversible inhibitors of Bruton’s tyrosine kinase reveals differences in on - and off - target inhibition HM71224, a selective Bruton’s tyrosine kinase inhibitor, attenuates the development of murine lupus, Biochim Biophys Acta Gen Subj 1864, 129531.

[39] Byrd, J. C., Harrington, B., O’Brien, S., Jones, J. A., Schuh, A., Devereux, S., Chaves, J., Wierda, W. G., Awan, F. T., Brown, J. R., Hillmen, P., Stephens, D. M., Ghia, P., Barrientos, J. C., Pagel, J. M., Woyach, J., Johnson, D., Huang, J., Wang, X., Kaptein, A., Lannutti, B. J., Covey, T., Fardis, M., McGreivy, J., Hamdy, A., Rothbaum, W., Izumi, R., Diacovo, T. G., Johnson, A. J., and Furman, R. R. (2016) Acalabrutinib (ACP-196) in Relapsed Chronic Lymphocytic Leukemia, N Engl J Med 374, 323–332.

[40] Caldwell, R. D., Qiu, H., Askew, B. C., Bender, A. T., Brugger, N., Camps, M., Dhanabal, M., Dutt, V., Eichhorn, T., Gardberg, A. S., Goutopoulos, A., Grenningloh, R., Head, J., Healey, B., Hodous, B. L., Huck, B. R., Johnson, T. L., Jones, C., Jones, R. C., Mochalkin, I., Morandi, F., Nguyen, N., Meyring, M., Potnick, J. R., Santos, D. C., Schmidt, R., Sherer, B., Shutes, A., Urbahns, K., Follis, A. V., Wegener, A. A., Zimmerli, S. C., and Liu-Bujalski, L. (2019) Discovery of Evobrutinib: An Oral, Potent, and Highly Selective, Covalent Bruton’s Tyrosine Kinase (BTK) Inhibitor for the Treatment of Immunological Diseases, J Med Chem 62, 7643–7655.

[41] Crawford, J. J., Johnson, A. R., Misner, D. L., Belmont, L. D., Castanedo, G., Choy, R., Coraggio, M., Dong, L., Eigenbrot, C., Erickson, R., Ghilardi, N., Hau, J., Katewa, A., Kohli, P. B., Lee, W., Lubach, J. W., McKenzie, B. S., Ortwine, D. F., Schutt, L., Tay, S., Wei, B., Reif, K., Liu, L., Wong, H., and Young, W. B. (2018) Discovery of GDC-0853: A Potent, Selective, and Noncovalent Bruton’s Tyrosine Kinase Inhibitor in Early Clinical Development, J Med Chem 61, 2227–2245.

[42] Dahl, K., Turner, T., Vasdev, N., Davis, M. I., Hunt, J. P., Herrgard, S., Ciceri, P., Wodicka, L. M., Pallares, G., Hocker, M., Treiber, D. K., Zarrinkar, P. P., and Davis, R. L. (2020) Radiosynthesis of a Bruton’s tyrosine kinase inhibitor, [(11) C]Tolebrutinib, via palladium-NiXantphos-mediated carbonylation Comprehensive analysis of kinase inhibitor selectivity Mechanism of Action and Target Identification: A Matter of Timing in Drug Discovery, J Labelled Comp Radiopharm 63, 482–487.

[43] Evans, E. K., Tester, R., Aslanian, S., Karp, R., Sheets, M., Labenski, M. T., Witowski, S. R., Lounsbury, H., Chaturvedi, P., Mazdiyasni, H., Zhu, Z., Nacht, M., Freed, M. I., Petter, R. C., Dubrovskiy, A., Singh, J., Westlin, W. F., Fabian, M. A., Biggs, W. H., 3rd, Treiber, D. K., Atteridge, C. E., Azimioara, M. D., Benedetti, M. G., Carter, T. A., Ciceri, P., Edeen, P. T., Floyd, M., Ford, J. M., Galvin, M., Gerlach, J. L., Grotzfeld, R. M., Herrgard, S., Insko, D. E., Insko, M. A., Lai, A. G., Lélias, J. M., Mehta, S. A., Milanov, Z. V., Velasco, A. M., Wodicka, L. M., Patel, H. K., Zarrinkar, P. P., Lockhart, D. J., Furman, R. R., Cheng, S., Lu, P., Setty, M., Perez, A. R., Guo, A., Racchumi, J., Xu, G., Wu, H., Ma, J., Steggerda, S. M., Coleman, M., Leslie, C., and Wang, Y. L. (2013) Inhibition of Btk with CC-292 provides early pharmacodynamic assessment of activity in mice and humans A small molecule-kinase interaction map for clinical kinase inhibitors Ibrutinib resistance in chronic lymphocytic leukemia, J Pharmacol Exp Ther 346, 219-228.

[44] García-Merino, A. (2021) Bruton’s Tyrosine Kinase Inhibitors: A New Generation of Promising Agents for Multiple Sclerosis Therapy, Cells 10.

[45] Goess, C., Harris, C. M., Murdock, S., McCarthy, R. W., Sampson, E., Twomey, R., Mathieu, S., Mario, R., Perham, M., Goedken, E. R., Long, A. J., Goldmann, L., Duan, R., Kragh, T., Wittmann, G., Weber, C., Lorenz, R., von Hundelshausen, P., Spannagl, M., and Siess, W. (2019) ABBV-105, a selective and irreversible inhibitor of Bruton’s tyrosine kinase, is efficacious in multiple preclinical models of inflammation Oral Bruton tyrosine kinase inhibitors block activation of the platelet Fc receptor CD32a (FcγRIIA): a new option in HIT?, Mod Rheumatol 29, 510–522.

[46] Guo, Y., Liu, Y., Hu, N., Yu, D., Zhou, C., Shi, G., Zhang, B., Wei, M., Liu, J., Luo, L., Tang, Z., Song, H., Guo, Y., Liu, X., Su, D., Zhang, S., Song, X., Zhou, X., Hong, Y., Chen, S., Cheng, Z., Young, S., Wei, Q., Wang, H., Wang, Q., Lv, L., Wang, F., Xu, H., Sun, H., Xing, H., Li, N., Zhang, W., Wang, Z., Liu, G., Sun, Z., Zhou, D., Li, W., Liu, L., Wang, L., Wang, Z., Angst, D., Gessier, F., Janser, P., Vulpetti, A., Wälchli, R., Beerli, C., Littlewood-Evans, A., Dawson, J., Nuesslein-Hildesheim, B., Wieczorek, G., Gutmann, S., Scheufler, C., Hinniger, A., Zimmerlin, A., Funhoff, E. G., Pulz, R., Cenni, B., Honigberg, L. A., Smith, A. M., Sirisawad, M., Verner, E., Loury, D., Chang, B., Li, S., Pan, Z., Thamm, D. H., Miller, R. A., and Buggy, J. J. (2019) Discovery of Zanubrutinib (BGB-3111), a Novel, Potent, and Selective Covalent Inhibitor of Bruton’s Tyrosine Kinase Discovery of LOU064 (Remibrutinib), a Potent and Highly Selective Covalent Inhibitor of Bruton’s Tyrosine Kinase The Bruton tyrosine kinase inhibitor PCI-32765 blocks B-cell activation and is efficacious in models of autoimmune disease and B-cell malignancy, J Med Chem 62, 7923-7940.

[47] Kim, Y. Y., Park, K. T., Jang, S. Y., Lee, K. H., Byun, J. Y., Suh, K. H., Lee, Y. M., Kim, Y. H., and Hwang, K. W. (2017) HM71224, a selective Bruton’s tyrosine kinase inhibitor, attenuates the development of murine lupus, Arthritis Res Ther 19, 211.

[48] Owens, T. D., Smith, P. F., Redfern, A., Xing, Y., Shu, J., Karr, D. E., Hartmann, S., Francesco, M. R., Langrish, C. L., Nunn, P. A., and Gourlay, S. G. (2022) Phase 1 clinical trial evaluating safety, exposure and pharmacodynamics of BTK inhibitor tolebrutinib (PRN2246, SAR442168), Clin Transl Sci 15, 442-450.

[49] Reiff, S. D., Mantel, R., Smith, L. L., Greene, J. T., Muhowski, E. M., Fabian, C. A., Goettl, V. M., Tran, M., Harrington, B. K., Rogers, K. A., Awan, F. T., Maddocks, K., Andritsos, L., Lehman, A. M., Sampath, D., Lapalombella, R., Eathiraj, S., Abbadessa, G., Schwartz, B., Johnson, A. J., Byrd, J. C., and Woyach, J. A. (2018) The BTK Inhibitor ARQ 531 Targets Ibrutinib-Resistant CLL and Richter Transformation, Cancer Discov 8, 1300–1315.

[50] Schafer, P. H., Kivitz, A. J., Ma, J., Korish, S., Sutherland, D., Li, L., Azaryan, A., Kosek, J., Adams, M., Capone, L., Hur, E. M., Hough, D. R., and Ringheim, G. E. (2020) Spebrutinib (CC-292) Affects Markers of B Cell Activation, Chemotaxis, and Osteoclasts in Patients with Rheumatoid Arthritis: Results from a Mechanistic Study, Rheumatol Ther 7, 101–119.

[51] Tam, C. S., Trotman, J., Opat, S., Burger, J. A., Cull, G., Gottlieb, D., Harrup, R., Johnston, P. B., Marlton, P., Munoz, J., Seymour, J. F., Simpson, D., Tedeschi, A., Elstrom, R., Yu, Y., Tang, Z., Han, L., Huang, J., Novotny, W., Wang, L., and Roberts, A. W. (2019) Phase 1 study of the selective BTK inhibitor zanubrutinib in B-cell malignancies and safety and efficacy evaluation in CLL, Blood 134, 851–859.

[52] Watterson, S. H., Liu, Q., Beaudoin Bertrand, M., Batt, D. G., Li, L., Pattoli, M. A., Skala, S., Cheng, L., Obermeier, M. T., Moore, R., Yang, Z., Vickery, R., Elzinga, P. A., Discenza, L., D’Arienzo, C., Gillooly, K. M., Taylor, T. L., Pulicicchio, C., Zhang, Y., Heimrich, E., McIntyre, K. W., Ruan, Q., Westhouse, R. A., Catlett, I. M., Zheng, N., Chaudhry, C., Dai, J., Galella, M. A., Tebben, A. J., Pokross, M., Li, J., Zhao, R., Smith, D., Rampulla, R., Allentoff, A., Wallace, M. A., Mathur, A., Salter-Cid, L., Macor, J. E., Carter, P. H., Fura, A., Burke, J. R., Tino, J. A., Caldwell, R. D., Qiu, H., Askew, B. C., Bender, A. T., Brugger, N., Camps, M., Dhanabal, M., Dutt, V., Eichhorn, T., Gardberg, A. S., Goutopoulos, A., Grenningloh, R., Head, J., Healey, B., Hodous, B. L., Huck, B. R., Johnson, T. L., Jones, C., Jones, R. C., Mochalkin, I., Morandi, F., Nguyen, N., Meyring, M., Potnick, J. R., Santos, D. C., Schmidt, R., Sherer, B., Shutes, A., Urbahns, K., Follis, A. V., Wegener, A. A., Zimmerli, S. C., Liu-Bujalski, L., Wen, T., Wang, J., Shi, Y., Qian, H., and Liu, P. (2019) Discovery of Branebrutinib (BMS-986195): A Strategy for Identifying a Highly Potent and Selective Covalent Inhibitor Providing Rapid in Vivo Inactivation of Bruton’s Tyrosine Kinase (BTK) Discovery of Evobrutinib: An Oral, Potent, and Highly Selective, Covalent Bruton’s Tyrosine Kinase (BTK) Inhibitor for the Treatment of Immunological Diseases Inhibitors targeting Bruton’s tyrosine kinase in cancers: drug development advances, J Med Chem 62, 3228–3250.

[53] Zhao, Y., Shu, Y., Lin, J., Chen, Z., Xie, Q., Bao, Y., Lu, L., Sun, N., Wang, Y., Watterson, S. H., Liu, Q., Beaudoin Bertrand, M., Batt, D. G., Li, L., Pattoli, M. A., Skala, S., Cheng, L., Obermeier, M. T., Moore, R., Yang, Z., Vickery, R., Elzinga, P. A., Discenza, L., D’Arienzo, C., Gillooly, K. M., Taylor, T. L., Pulicicchio, C., Zhang, Y., Heimrich, E., McIntyre, K. W., Ruan, Q., Westhouse, R. A., Catlett, I. M., Zheng, N., Chaudhry, C., Dai, J., Galella, M. A., Tebben, A. J., Pokross, M., Li, J., Zhao, R., Smith, D., Rampulla, R., Allentoff, A., Wallace, M. A., Mathur, A., Salter-Cid, L., Macor, J. E., Carter, P. H., Fura, A., Burke, J. R., Tino, J. A., Caldwell, R. D., Qiu, H., Askew, B. C., Bender, A. T., Brugger, N., Camps, M., Dhanabal, M., Dutt, V., Eichhorn, T., Gardberg, A. S., Goutopoulos, A., Grenningloh, R., Head, J., Healey, B., Hodous, B. L., Huck, B. R., Johnson, T. L., Jones, C., Jones, R. C., Mochalkin, I., Morandi, F., Nguyen, N., Meyring, M., Potnick, J. R., Santos, D. C., Schmidt, R., Sherer, B., Shutes, A., Urbahns, K., Follis, A. V., Wegener, A. A., Zimmerli, S. C., Liu-Bujalski, L., Wen, T., Wang, J., Shi, Y., Qian, H., and Liu, P. (2021) Discovery of novel BTK PROTACs for B-Cell lymphomas Discovery of Branebrutinib (BMS-986195): A Strategy for Identifying a Highly Potent and Selective Covalent Inhibitor Providing Rapid in Vivo Inactivation of Bruton’s Tyrosine Kinase (BTK) Discovery of Evobrutinib: An Oral, Potent, and Highly Selective, Covalent Bruton’s Tyrosine Kinase (BTK) Inhibitor for the Treatment of Immunological Diseases Inhibitors targeting Bruton’s tyrosine kinase in cancers: drug development advances, Eur J Med Chem 225, 113820.

[54] Wu, J., Zhang, M., and Liu, D. (2016) Acalabrutinib (ACP-196): a selective second-generation BTK inhibitor, J Hematol Oncol 9, 21.

[55] Berg, E. L. (2017) Phenotypic chemical biology for predicting safety and efficacy, Drug Discov Today Technol 23, 53–60.

[56] Davis, R. L. (2020) Mechanism of Action and Target Identification: A Matter of Timing in Drug Discovery, iScience 23, 101487.

[57] Lin, A., Giuliano, C. J., Palladino, A., John, K. M., Abramowicz, C., Yuan, M. L., Sausville, E. L., Lukow, D. A., Liu, L., Chait, A. R., Galluzzo, Z. C., Tucker, C., and Sheltzer, J. M. (2019) Off-target toxicity is a common mechanism of action of cancer drugs undergoing clinical trials, Sci Transl Med 11.

[58] Isenberg, D., Furie, R., Jones, N. S., Guibord, P., Galanter, J., Lee, C., McGregor, A., Toth, B., Rae, J., Hwang, O., Desai, R., Lokku, A., Ramamoorthi, N., Hackney, J. A., Miranda, P., de Souza, V. A., Jaller-Raad, J. J., Maura Fernandes, A., Garcia Salinas, R., Chinn, L. W., Townsend, M. J., Morimoto, A. M., Tuckwell, K., Jarmoskaite, I., AlSadhan, I., Vaidyanathan, P. P., and Herschlag, D. (2021) Efficacy, Safety, and Pharmacodynamic Effects of the Bruton’s Tyrosine Kinase Inhibitor Fenebrutinib (GDC-0853) in Systemic Lupus Erythematosus: Results of a Phase II, Randomized, Double-Blind, Placebo-Controlled Trial How to measure and evaluate binding affinities, Arthritis Rheumatol 73, 1835-1846.

[59] Brown, J. R., Eichhorst, B., Hillmen, P., Jurczak, W., Kaźmierczak, M., Lamanna, N., O’Brien, S. M., Tam, C. S., Qiu, L., Zhou, K., Simkovic, M., Mayer, J., Gillespie-Twardy, A., Ferrajoli, A., Ganly, P. S., Weinkove, R., Grosicki, S., Mital, A., Robak, T., Osterborg, A., Yimer, H. A., Salmi, T., Wang, M. D., Fu, L., Li, J., Wu, K., Cohen, A., and Shadman, M. (2023) Zanubrutinib or Ibrutinib in Relapsed or Refractory Chronic Lymphocytic Leukemia, N Engl J Med 388, 319–332.

[60] Byrd, J. C., Hillmen, P., Ghia, P., Kater, A. P., Chanan-Khan, A., Furman, R. R., O’Brien, S., Yenerel, M. N., Illés, A., Kay, N., Garcia-Marco, J. A., Mato, A., Pinilla-Ibarz, J., Seymour, J. F., Lepretre, S., Stilgenbauer, S., Robak, T., Rothbaum, W., Izumi, R., Hamdy, A., Patel, P., Higgins, K., Sohoni, S., and Jurczak, W. (2021) Acalabrutinib Versus Ibrutinib in Previously Treated Chronic Lymphocytic Leukemia: Results of the First Randomized Phase III Trial, J Clin Oncol 39, 3441–3452.

[61] Wang, X., Wong, J., Sevinsky, C. J., Kokabee, L., Khan, F., Sun, Y., and Conklin, D. S. (2016) Bruton’s Tyrosine Kinase Inhibitors Prevent Therapeutic Escape in Breast Cancer Cells, Mol Cancer Ther 15, 2198–2208.

[62] Wu, H., Hu, C., Wang, A., Weisberg, E. L., Chen, Y., Yun, C. H., Wang, W., Liu, Y., Liu, X., Tian, B., Wang, J., Zhao, Z., Liang, Y., Li, B., Wang, L., Wang, B., Chen, C., Buhrlage, S. J., Qi, Z., Zou, F., Nonami, A., Li, Y., Fernandes, S. M., Adamia, S., Stone, R. M., Galinsky, I. A., Wang, X., Yang, G., Griffin, J. D., Brown, J. R., Eck, M. J., Liu, J., Gray, N. S., and Liu, Q. (2016) Discovery of a BTK/MNK dual inhibitor for lymphoma and leukemia, Leukemia 30, 173–181.

[63] Melton, A. C., Melrose, J., Alajoki, L., Privat, S., Cho, H., Brown, N., Plavec, A. M., Nguyen, D., Johnston, E. D., Yang, J., Polokoff, M. A., Plavec, I., Berg, E. L., and O’Mahony, A. (2013) Regulation of IL-17A production is distinct from IL-17F in a primary human cell co-culture model of T cell-mediated B cell activation, PLoS One 8, e58966.

[64] Luo, Q. Y., Zhou, S. N., Pan, W. T., Sun, J., Yang, L. Q., Zhang, L., Qiu, M. Z., Yang, D. J., Long, M., Beckwith, K., Do, P., Mundy, B. L., Gordon, A., Lehman, A. M., Maddocks, K. J., Cheney, C., Jones, J. A., Flynn, J. M., Andritsos, L. A., Awan, F., Fraietta, J. A., June, C. H., Maus, M. V., Woyach, J. A., Caligiuri, M. A., Johnson, A. J., Muthusamy, N., and Byrd, J. C. (2021) A multi-kinase inhibitor APG-2449 enhances the antitumor effect of ibrutinib in esophageal squamous cell carcinoma via EGFR/FAK pathway inhibition Ibrutinib treatment improves T cell number and function in CLL patients, Biochem Pharmacol 183, 114318.

[65] Tan, B., Huang, Y., Zhang, B., Lin, N., Pal Singh, S., Dammeijer, F., and Hendriks, R. W. (2020) The effect of ibrutinib on radiosensitivity in pancreatic cancer cells by targeting EGFR/AKT/mTOR signaling pathway Correction to: Role of Bruton’s tyrosine kinase in B cells and malignancies, Biomed Pharmacother 128, 110133.

[66] Di Paolo, J. A., Huang, T., Balazs, M., Barbosa, J., Barck, K. H., Bravo, B. J., Carano, R. A., Darrow, J., Davies, D. R., DeForge, L. E., Diehl, L., Ferrando, R., Gallion, S. L., Giannetti, A. M., Gribling, P., Hurez, V., Hymowitz, S. G., Jones, R., Kropf, J. E., Lee, W. P., Maciejewski, P. M., Mitchell, S. A., Rong, H., Staker, B. L., Whitney, J. A., Yeh, S., Young, W. B., Yu, C., Zhang, J., Reif, K., and Currie, K. S. (2011) Specific Btk inhibition suppresses B cell- and myeloid cell-mediated arthritis, Nat Chem Biol 7, 41–50.

[67] Cohen, S., Tuckwell, K., Katsumoto, T. R., Zhao, R., Galanter, J., Lee, C., Rae, J., Toth, B., Ramamoorthi, N., Hackney, J. A., Berman, A., Damjanov, N., Fedkov, D., Jeka, S., Chinn, L. W., Townsend, M. J., Morimoto, A. M., and Genovese, M. C. (2020) Fenebrutinib versus Placebo or Adalimumab in Rheumatoid Arthritis: A Randomized, Double-Blind, Phase II Trial (ANDES Study), Arthritis Rheumatol 72, 1435–1446.

68. [68] Goldmann, L., Duan, R., Kragh, T., Wittmann, G., Weber, C., Lorenz, R., von Hundelshausen, P., Spannagl, M., and Siess, W. (2019) Oral Bruton tyrosine kinase inhibitors block activation of the platelet Fc receptor CD32a (FcγRIIA): a new option in HIT?, Blood Adv 3, 4021–4033.

[69] Müller, C. W., Rey, F. A., Sodeoka, M., Verdine, G. L., Harrison, S. C., Maddocks, K. J., Ruppert, A. S., Lozanski, G., Heerema, N. A., Zhao, W., Abruzzo, L., Lozanski, A., Davis, M., Gordon, A., Smith, L. L., Mantel, R., Jones, J. A., Flynn, J. M., Jaglowski, S. M., Andritsos, L. A., Awan, F., Blum, K. A., Grever, M. R., Johnson, A. J., Byrd, J. C., and Woyach, J. A. (1995) Structure of the NF-kappa B p50 homodimer bound to DNA Etiology of Ibrutinib Therapy Discontinuation and Outcomes in Patients With Chronic Lymphocytic Leukemia, Nature 373, 311–317.

[70] Davis, M. I., Hunt, J. P., Herrgard, S., Ciceri, P., Wodicka, L. M., Pallares, G., Hocker, M., Treiber, D. K., Zarrinkar, P. P., and Davis, R. L. (2011) Comprehensive analysis of kinase inhibitor selectivity Mechanism of Action and Target Identification: A Matter of Timing in Drug Discovery, Nat Biotechnol 29, 1046–1051.

[71] Fabian, M. A., Biggs, W. H., 3rd, Treiber, D. K., Atteridge, C. E., Azimioara, M. D., Benedetti, M. G., Carter, T. A., Ciceri, P., Edeen, P. T., Floyd, M., Ford, J. M., Galvin, M., Gerlach, J. L., Grotzfeld, R. M., Herrgard, S., Insko, D. E., Insko, M. A., Lai, A. G., Lélias, J. M., Mehta, S. A., Milanov, Z. V., Velasco, A. M., Wodicka, L. M., Patel, H. K., Zarrinkar, P. P., Lockhart, D. J., Furman, R. R., Cheng, S., Lu, P., Setty, M., Perez, A. R., Guo, A., Racchumi, J., Xu, G., Wu, H., Ma, J., Steggerda, S. M., Coleman, M., Leslie, C., and Wang, Y. L. (2005) A small molecule-kinase interaction map for clinical kinase inhibitors Ibrutinib resistance in chronic lymphocytic leukemia, Nat Biotechnol 23, 329-336.

[72] Wodicka, L. M., Ciceri, P., Davis, M. I., Hunt, J. P., Floyd, M., Salerno, S., Hua, X. H., Ford, J. M., Armstrong, R. C., Zarrinkar, P. P., Treiber, D. K., Wang, M. L., Rule, S., Martin, P., Goy, A., Auer, R., Kahl, B. S., Jurczak, W., Advani, R. H., Romaguera, J. E., Williams, M. E., Barrientos, J. C., Chmielowska, E., Radford, J., Stilgenbauer, S., Dreyling, M., Jedrzejczak, W. W., Johnson, P., Spurgeon, S. E., Li, L., Zhang, L., Newberry, K., Ou, Z., Cheng, N., Fang, B., McGreivy, J., Clow, F., Buggy, J. J., Chang, B. Y., Beaupre, D. M., Kunkel, L. A., and Blum, K. A. (2010) Activation state-dependent binding of small molecule kinase inhibitors: structural insights from biochemistry Targeting BTK with ibrutinib in relapsed or refractory mantle-cell lymphoma, Chem Biol 17, 1241–1249.

[73] Shah, F., Stepan, A. F., O’Mahony, A., Velichko, S., Folias, A. E., Houle, C., Shaffer, C. L., Marcek, J., Whritenour, J., Stanton, R., Berg, E. L., Schafer, P. H., Kivitz, A. J., Ma, J., Korish, S., Sutherland, D., Li, L., Azaryan, A., Kosek, J., Adams, M., Capone, L., Hur, E. M., Hough, D. R., and Ringheim, G. E. (2017) Mechanisms of Skin Toxicity Associated with Metabotropic Glutamate Receptor 5 Negative Allosteric Modulators Spebrutinib (CC- 292) Affects Markers of B Cell Activation, Chemotaxis, and Osteoclasts in Patients with Rheumatoid Arthritis: Results from a Mechanistic Study, Cell Chem Biol 24, 858–869 e855.

