## Supplemental Material for "Comprehensive Characterization of BTK Inhibitor Specificity, Potency, and Biological Effects: Insights into Covalent and Non-covalent Mechanistic Signatures"

| Compound Name | Mechanism | BTK binding POC | TEC binding POC | BMX binding POC | TXK binding POC | ITK binding POC |
|---|---|---|---|---|---|---|
| Ibrutinib | covalent | 0 | 5.9 | 6.4 | 0.5 | 3.6 |
| Poseltinib | non-covalent | 0.1 | 8.5 | 12 | 4.4 | 1 |
| Branebrutinib | covalent | 0 | 7.8 | 11 | 2 | 17 |
| Spebrutinib | covalent | 0 | 8.1 | 10 | 4.5 | 51 |
| Tolebrutinib | covalent | 0 | 4.2 | 11 | 1.1 | 67 |
| Zanubrutinib | covalent | 0 | 8.7 | 10 | 2.2 | 74 |
| Nemtabrutinib | non-covalent | 0.6 | 7.3 | 8.5 | 2.7 | 86 |
| Fenebrutinib | non-covalent | 0 | 17 | 16 | 53 | 25 |
| Tirabrutinib | covalent | 0.35 | 9.4 | 16 | 14 | 100 |
| Evobrutinib | covalent | 0.65 | 10 | 13 | 23 | 95 |
| Elsubrutinib | covalent | 0.1 | 18 | 20 | 25 | 94 |
| Orelabrutinib | covalent | 0.35 | 12 | 27 | 34 | 100 |
| Pirtobrutinib | non-covalent | 0 | 12 | 51 | 11 | 100 |
| Acalabrutinib | covalent | 0.2 | 14 | 52 | 47 | 100 |
| Remibrutinib | covalent | 0.15 | 20 | 47 | 97 | 97 |

**Supplemental Table 1. Comparison of BTK inhibitor specificity to TEC family members.** The percent of control (POC) is the percent of kinase that was competed off its control ligand relative to the positive and negative controls at the screening concentration (Karaman et al. 2008). A zero POC signifies that the compound competition was comparable to that of the positive control. TEC family members are Burton's tyrosine kinase (BTK), tyrosine kinase expressed in hepatocellular carcinoma (TEC), bone marrow expressed kinase (BMX/ETK), and T cell expressed kinase (TXK/RLK), and interleukin-2-inducible T cell kinase (ITK/TSK).

| Compound Name | Mechanism | BLK binding POC | EGFR binding POC | ERBB2 binding POC | ERBB3 binding POC | ERBB4 binding POC | JAK3 binding POC | MKK7 binding POC |
|---|---|---|---|---|---|---|---|---|
| Ibrutinib | covalent | 0.25 | 0 | 0.1 | 0.15 | 0 | 0.7 | 0.15 |
| Tolebrutinib | covalent | 0.05 | 0 | 0 | 0.8 | 0 | 36 | 19 |
| Nemtabrutinib | non-covalent | 0.3 | 0 | 0 | 0.9 | 0.35 | 86 | 39 |
| Zanubrutinib | covalent | 0.3 | 0 | 7.3 | 23 | 0.75 | 20 | 55 |
| Poseltinib | non-covalent | 0.85 | 1.2 | 8.7 | 97 | 0.15 | 0.45 | 100 |
| Pirtobrutinib | non-covalent | 69 | 0.5 | 5.9 | 97 | 2 | 77 | 91 |
| Acalabrutinib | covalent | 57 | 61 | 4.7 | 64 | 6.9 | 80 | 75 |
| Spebrutinib | covalent | 22 | 49 | 57 | 83 | 17 | 0 | 79 |
| Elsubrutinib | covalent | 15 | 90 | 73 | 74 | 91 | 0.15 | 24 |
| Tirabrutinib | covalent | 14 | 74 | 26 | 12 | 85 | 73 | 87 |
| Branebrutinib | covalent | 3.2 | 77 | 100 | 75 | 75 | 48 | 100 |
| Evobrutinib | covalent | 22 | 88 | 91 | 97 | 32 | 100 | 100 |
| Orelabrutinib | covalent | 52 | 68 | 100 | 88 | 96 | 89 | 100 |
| Fenebrutinib | non-covalent | 60 | 95 | 97 | 78 | 100 | 95 | 89 |
| Remibrutinib | covalent | 100 | 99 | 100 | 100 | 100 | 100 | 100 |

**Supplemental Table 2. Comparison of BTK inhibitor specificity to HER2+ associated kinases.** The percent of control (POC) is the percent of kinase that was competed off its control ligand relative to the positive and negative controls at the screening concentration (Karaman et al. 2008). A zero POC signifies that the compound competition was comparable to that of the positive control. Abbreviations: B lymphocyte kinase (BLK), epidermal growth factor receptor (EGFR), human epidermal growth factor receptor 2 (HER2), Erb-B2 receptor tyrosine kinases 2, 3, and 4 (ERBB2, ERBB3, and ERBB4, respectively), Janus kinase 3 (JAK3), and mitogen-activated protein kinase kinase 7 (MKK7). Note that the JAK3 POC is for its JH1 catalytic domain.

| Compound Name | Mechanism | Average in-house Kd for BTK (nM) (number of replicates) | Average published IC50 or Kd for BTK (nM) (number of replicates) | Fold difference between in-house and published IC50 or Kd |
|---|---|---|---|---|
| Acalabrutinib | covalent | 4.2 (n=7) | 7.4 (n=5) | 1.78 |
| Branebrutinib | covalent | 0.11 (n=7) | 0.1 (n=1) | 0.91 |
| Elsubrutinib | covalent | 2.9 (n=7) | 180 (n=1) | 62.07 |
| Evobrutinib | covalent | 18 (n=7) | 19 (n=3) | 1.06 |
| Ibrutinib | covalent | 0.62 (n=7) | 0.83 (n=10) | 1.35 |
| Orelabrutinib | covalent | 4.6 (n=7) | 1.6 (n=1) | 0.35 |
| Remibrutinib | covalent | 0.25 (n=7) | 1.3 (n=1) | 5.2 |
| Spebrutinib | covalent | 2.3 (n=7) | 5.2 (n=5) | 2.29 |
| Tirabrutinib | covalent | 5.6 (n=7) | 9.4 (n=3) | 1.69 |
| Tolebrutinib | covalent | 1.1 (n=7) | 0.55 (n=2) | 0.49 |
| Zanubrutinib | covalent | 0.38 (n=7) | 0.34 (n=3) | 0.89 |
| Fenebrutinib | non-covalent | 1.5 (n=1) | 1.3 (n=2) | 0.87 |
| Pirtobrutinib | non-covalent | 0.81 (n=1) | 2 (n=2) | 2.47 |
| Nemtabrutinib | non-covalent | 9.9 (n=1) | 31 (n=3) | 3.13 |
| Poseltinib | non-covalent | 1.5 (n=1) | 3 (n=2) | 2 |

**Supplemental Table 3. Comparison of in-house and published BTK inhibitor binding affinity to BTK.** The binding mechanism to wild-type BTK is listed in the second column. The number of replicates for BTK IC50(s) or Kd(s) is shown in parentheses after the average IC50 or Kd value. The fold difference between in-house and published results is shown in the fifth column.

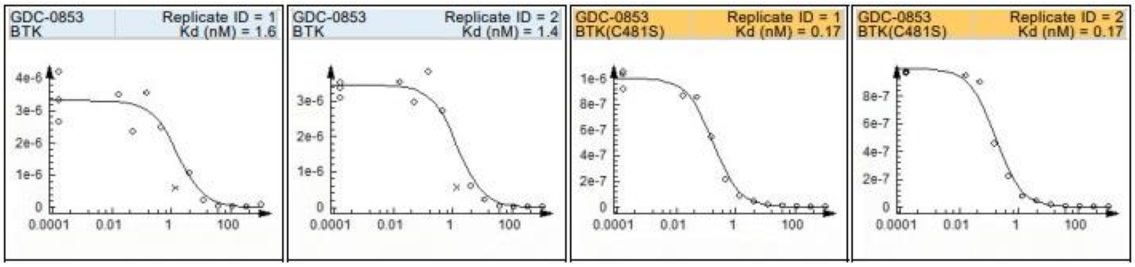

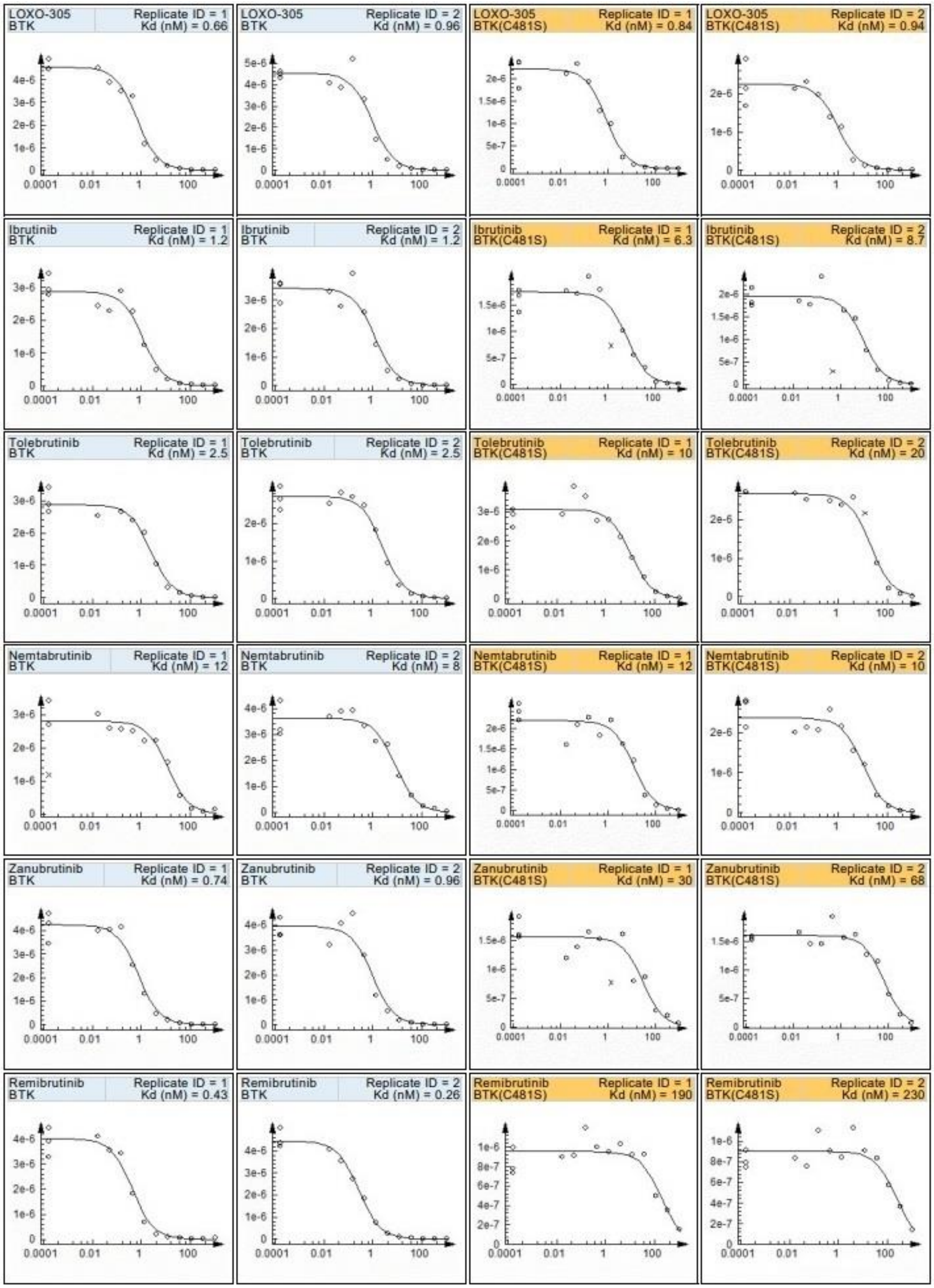
LOXO-305 BTK Replicate ID = 1 Kd (nM) = 0.66
LOXO-305 BTK Replicate ID = 2 Kd (nM) = 0.96
LOXO-305 BTK(C481S) Replicate ID = 1 Kd (nM) = 0.84
LOXO-305 BTK(C481S) Replicate ID = 2 Kd (nM) = 0.94
Ibrutinib BTK Replicate ID = 1 Kd (nM) = 1.2
Ibrutinib BTK Replicate ID = 2 Kd (nM) = 1.2
Ibrutinib BTK(C481S) Replicate ID = 1 Kd (nM) = 6.3
Ibrutinib BTK(C481S) Replicate ID = 2 Kd (nM) = 8.7
Tolebrutinib BTK Replicate ID = 1 Kd (nM) = 2.5
Tolebrutinib BTK Replicate ID = 2 Kd (nM) = 2.5
Tolebrutinib BTK(C481S) Replicate ID = 1 Kd (nM) = 10
Tolebrutinib BTK(C481S) Replicate ID = 2 Kd (nM) = 20
Nemtabrutinib BTK Replicate ID = 1 Kd (nM) = 12
Nemtabrutinib BTK Replicate ID = 2 Kd (nM) = 8
Nemtabrutinib BTK(C481S) Replicate ID = 1 Kd (nM) = 12
Nemtabrutinib BTK(C481S) Replicate ID = 2 Kd (nM) = 10
Zanubrutinib BTK Replicate ID = 1 Kd (nM) = 0.74
Zanubrutinib BTK Replicate ID = 2 Kd (nM) = 0.96
Zanubrutinib BTK(C481S) Replicate ID = 1 Kd (nM) = 30
Zanubrutinib BTK(C481S) Replicate ID = 2 Kd (nM) = 68
Remibrutinib BTK Replicate ID = 1 Kd (nM) = 0.43
Remibrutinib BTK Replicate ID = 2 Kd (nM) = 0.26
Remibrutinib BTK(C481S) Replicate ID = 1 Kd (nM) = 190
Remibrutinib BTK(C481S) Replicate ID = 2 Kd (nM) = 230
0.0001
0.01
1
100

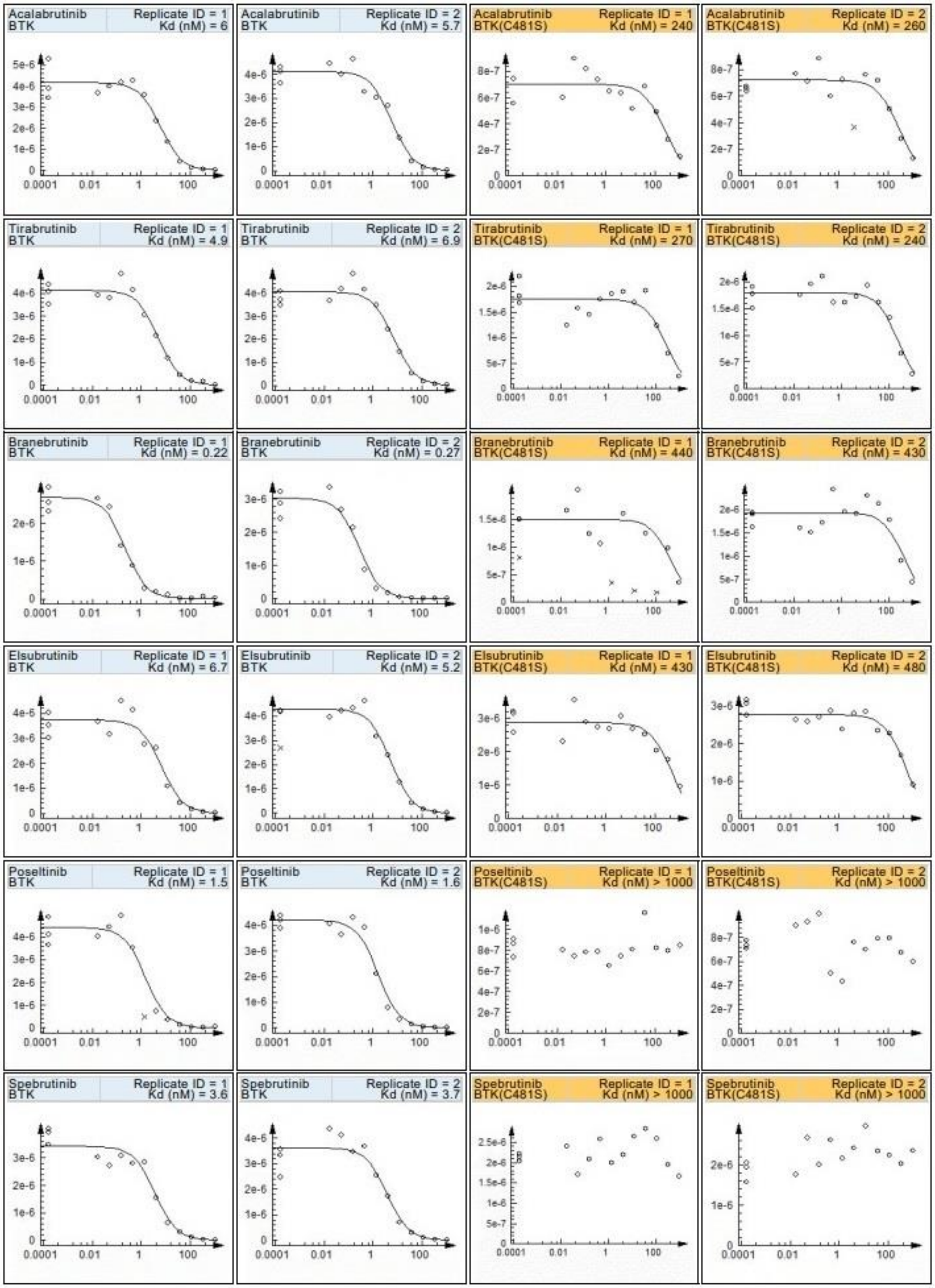
Acalabrutinib BTK Replicate ID = 1 Kd (nM) = 6
Acalabrutinib BTK Replicate ID = 2 Kd (nM) = 5.7
Acalabrutinib BTK(C481S) Replicate ID = 1 Kd (nM) = 240
Acalabrutinib BTK(C481S) Replicate ID = 2 Kd (nM) = 260
Tirabrutinib BTK Replicate ID = 1 Kd (nM) = 4.9
Tirabrutinib BTK Replicate ID = 2 Kd (nM) = 6.9
Tirabrutinib BTK(C481S) Replicate ID = 1 Kd (nM) = 270
Tirabrutinib BTK(C481S) Replicate ID = 2 Kd (nM) = 240
Branebrutinib BTK Replicate ID = 1 Kd (nM) = 0.22
Branebrutinib BTK Replicate ID = 2 Kd (nM) = 0.27
Branebrutinib BTK(C481S) Replicate ID = 1 Kd (nM) = 440
Branebrutinib BTK(C481S) Replicate ID = 2 Kd (nM) = 430
Elsubrutinib BTK Replicate ID = 1 Kd (nM) = 6.7
Elsubrutinib BTK Replicate ID = 2 Kd (nM) = 5.2
Elsubrutinib BTK(C481S) Replicate ID = 1 Kd (nM) = 430
Elsubrutinib BTK(C481S) Replicate ID = 2 Kd (nM) = 480
Poseltinib BTK Replicate ID = 1 Kd (nM) = 1.5
Poseltinib BTK Replicate ID = 2 Kd (nM) = 1.6
Poseltinib BTK(C481S) Replicate ID = 1 Kd (nM) > 1000
Poseltinib BTK(C481S) Replicate ID = 2 Kd (nM) > 1000
Spebrutinib BTK Replicate ID = 1 Kd (nM) = 3.6
Spebrutinib BTK Replicate ID = 2 Kd (nM) = 3.7
Spebrutinib BTK(C481S) Replicate ID = 1 Kd (nM) > 1000
Spebrutinib BTK(C481S) Replicate ID = 2 Kd (nM) > 1000

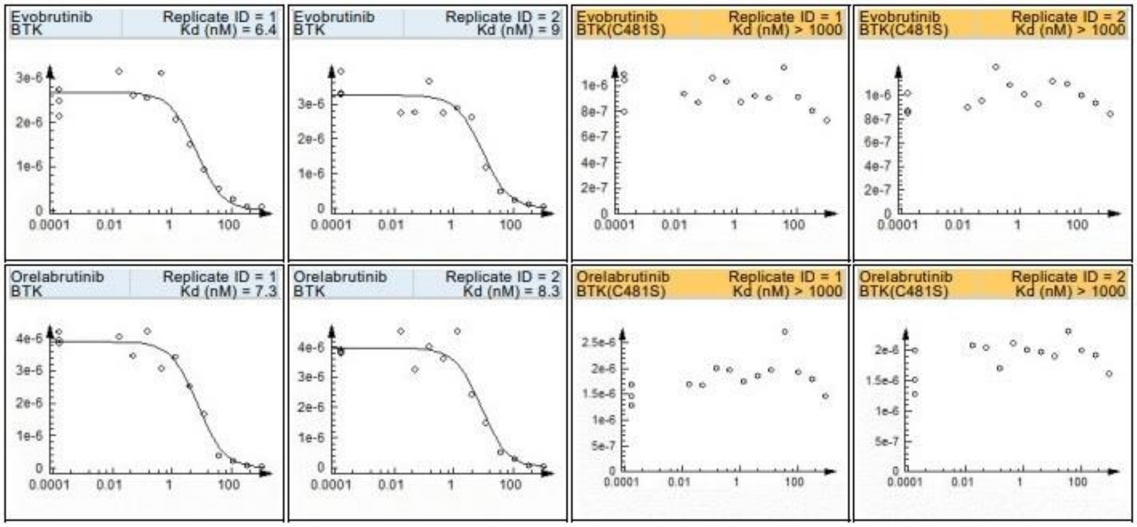


**Supplemental Figure 1. Potency curves of 15 BTK inhibitors to BTK and mutant BTK (BTK C481S**). 11-point dose response curves (or lack thereof) for each BTK inhibitor to wild-type BTK and a BTK mutant (BTK C481S). These curves were used to determine the binding affinities (Kds) of these interactions. The BTK inhibitors are placed in order of most potent to both BTK and BTK C481S (top) to least potent (bottom). Outlier data points marked with an "x" were not used for Kd determination. Each y-axis scales with the highest measured signal for that replicate. The x-axis is in units of nanomolar.
